# The DDIT4L-TOM40-ATP5A pathway suppresses glioblastoma oncogenesis

**DOI:** 10.1101/2025.07.27.666981

**Authors:** Pu You, Zeng-Xin Qi, Hao Luo, Bing Cai, Xu-Ye Hu, Jing Pan, Pan-Pan Gan, Xin-Ying Zeng, Ri-Jing Liao, Qisheng Tang, Ye Yu, Zhen-Ning Zhang, Bojie Yang, Liang Chen, Xu Zhang, Kai-Cheng Li, Ying Mao

**Affiliations:** Department of Neurosurgery, Huashan Hospital, Shanghai Medical College, State Key Laboratory of Medical Neurobiology, MOE Frontiers Center for Brain Science and Institutes of Brain Science, Fudan University, Shanghai 200040, China; Institute of Brain-Intelligence Science and Technology, Zhangjiang Laboratory, Shanghai 200031, China; Shanghai Clinical Research Center, Chinese Academy of Sciences/XuHui Central Hospital, Shanghai 200031, China; School of Life Science and Technology, ShanghaiTech University, Shanghai 201210, China; School of Traditional Chinese Pharmacy, China Pharmaceutical University, Nanjing 210009, China; School of Life Sciences and Technology, Tongji University, Shanghai 200092, China.; Tianqiao and Chrissy Chen Institute Clinical Translational Research Center, Shanghai 200040, China; Shanghai QuietD Biotechnology Co., Ltd., Shanghai 201210, China

**Keywords:** DDIT4L, TOM40, GBM oncogenesis, Mitochondrial dynamics, synthetic peptide

## Abstract

The characteristics of glioblastoma (GBM), including resistance to cell death and aberrant energy metabolism, are associated with the function of mitochondria. To date, the inherent suppressors that couple mitochondrial function with cell death remain unclear. DNA damage-inducible transcript 4-like (DDIT4L) expression in human gliomas and its association with patient survival was determined by public databases, western-blot and immunostaining. The role of DDIT4L in regulating tumor growth was analyzed using lentivirus and PDX model. The mechanism of DDIT4L on GBM mitochondrial function was determined by different assays. Here, we show that DDIT4L is an endogenous inhibitor of GBM via suppressing mitochondrial function. DDIT4L was expressed at high levels in GBM and transported into mitochondria via single import channel TOM40 other than the import receptors TOM22, TOM20 or TOM70. Then, DDIT4L interacted with the α subunit of ATP synthase, inhibited mitochondrial function and induced tumor cell apoptosis. Furthermore, the synthetic peptide DDIT4L^V125-P132^ suppressed GBM oncogenesis. Based on these results, DDIT4L suppresses GBM oncogenesis through both TOM40- and ATP5A-dependent mechanisms to curb mitochondrial dynamics and induce cell apoptosis, and the DDIT4L^V125-P132^ peptide is a potential therapy for GBM.

## Introduction

The translocase of the outer mitochondrial membrane (TOM complex) is the main entry point for nuclear-encoded mitochondrial proteins into mitochondria. In mammals the TOM complex consists of several components, including the receptor proteins TOM70, TOM22 or TOM20, and the protein-conducting channel protein TOM40, as well as the three smaller proteins, TOM7, TOM6 and TOM5^1^. Translocation of cytosolic proteins into mitochondria usually requires the N-terminal signal sequences which are 20-50 amino acids in length and usually contain amphipathic helices with Arg and Lys residues on one side and hydrophobic residues on the other. The interaction between cytosol proteins with import receptor TOM20, TOM22 or TOM70 is required for all four pathways that translocate these cytosol proteins into the mitochondria^2^. Subsequently, the cytosol proteins are transferred from import receptors to the general import channel, TOM40. Therefore, TOM40 is considered as transport pore other than the recognition import receptor in the translocation pathways. Theoretically, a protein will not be translocated into the mitochondrial matrix without interacting with the import receptors TOM20, TOM22 or TOM70. In the present study, we found that DDIT4L does not bind out-of-membrane receptors and inhibits mitochondrial function by targeting TOM40 in GBM cells. Under normal conditions, the TOM complex consists of two TOM40 proteins, two TOM22 proteins and two TOM5 proteins^3^. In GBM, TOM40 is expressed at higher levels than TOM22 and TOM5 according to the CGGA database. We unexpectedly observed that DDIT4L can be transported into mitochondrial matrix via single TOM40 other than TOM complex. Therefore, TOM40 may be a general import pore functioning as a single molecule and not as a TOM complex in GBM, which provides an opportunity for cytoplasmic proteins to interact with TOM40 and regulate mitochondrial function. In this study, we found that TOM40 binds and transports DDIT4L into the inner membrane of mitochondria, inhibits mitochondrial function and elicits cellular apoptosis, leading to the suppression of GBM oncogenesis.

The DDIT4L gene was first identified from a genetic screen for negative regulators of the *Drosophila* target of rapamycin (TOR) pathway, termed *Charybdis*^4^. In mammals, it is named DDIT4L (a.k.a. REDD2/RTP801L); while its homolog DDIT4 (also called REDD1/RTP801) shares ∼36% amino acid sequence homology with DDIT4L. These two proteins are upstream inhibitors of mechanistic TOR (mTOR) in several tissues and cell models^5-7^. In human tissues, DDIT4L is mainly expressed in skeletal muscle, while DDIT4 is expressed in almost all tissues^8^. *DDIT4L* was found to promote autophagy and inhibit pathological cardiac hypertrophy^6^. In the present study, we found that DDIT4L directly binds to the α subunit of ATP synthase via TOM40 translocation to inhibit mitochondrial function in GBM cells. Mitochondrial ATP synthase generates most cellular ATP under aerobic conditions. As we all known, the ATP synthase has two components: F_1_ and F_0_. The F_1_ portion, including subunits α (ATP5A) and β (ATP5B), catalyzes ATP synthesis, while the F_0_ portion forms a channel to translocate protons according to their gradient^9-11^. Previous studies have shown that D2HG produced in IDH mutant-type glioma binds ATP5B, inhibiting tumor growth^12,13^. In this study, we found that DDIT4L directly binds ATP5A to inhibit the activity of ATP synthase and elicit cellular apoptosis, leading to the suppression of GBM oncogenesis.

## Materials and Methods

### Human Tumor and cell Cultures and PDX model

Human tumor tissue collection and PDX model generation was conducted with the approval of the Huashan Hospital of Fudan University and the Zhangjiang Laboratory Institutional Review Board (SHIRB, KY2015-256). Informed consent was obtained from all subjects involved in this study. GBM and LGG tissues were obtained from newly diagnosed patients with the following clinical characteristics: hGBM21359, 71 year old male; hGBM11879, 68 year old male; hGBM11271, 42 year old male; hGBM12913, 64 year old female; hGBM79150, 53 year old female; hGBM21686, 28 year old female; hLGG14023, 38 year old male, grade III; hLGG11002, 53 year old male, grade III; hL21845, 56 year old male, grade III; hLGG12101, 39 year old female, grade II; hLGG21823, 24 year old male, grade II.

Patient-derived primary GBM cells from hGBM11879 and hGBM11271 were maintained in neurosphere medium (DMEM/F12, ThermoFisher, 11330032), supplemented with B27 (0.5X, ThermoFisher, 17504044), N2(0.5X, ThermoFisher, 17504048), bFGF (20 ng/ml, Peprotech, 100-18B) and EGF (20 ng/ml, Peprotech, 100-47), BSA (0.5mg/ml, Sigma-Aldrich, A9418).

Intracranial transplantation of primary GBM cells to establish tumor xenografts was performed as described^14^. Briefly, 48 hr after lentiviral infection, 2×10^5^ primary GBM cells were implanted into the left CPU region (from bregma, cells were injected 2.0 mm lateral, 0 mm caudal, 2.75 mm ventral) of male Nod-SCID or Nude mice. The 8-week old male littermates were used for transplantation and randomized to receive primary GBM cells expressing DDIT4L or control vector. For the survival experiments, animals were maintained until manifestation of neurological signs or for 120 days post-transplantation. To monitor tumor growth, mice transplanted with primary GBM cells, which stably expressed firefly luciferase, were monitored by bioluminescence imaging longitudinally using the IVIS100 bioluminescence imaging system. To analyze the effect of peptide on the growth of GBM cells in vivo, osmotic pumps (ALZET®, 1002) were implanted subcutaneously to direct deliver TAT-DDIT4L^V125-P132^ peptide or scrambling peptide (dissolved in PBS) to the injection site via brain infusion kit (ALZET®) a week after intracranial tumor cell injection.

### Immunostaining

Frozen brains were sectioned to 8-15 μm thickness. Sections were blocked in 10% normal donkey serum prepared in PBS and 0.3% TritonX-100 for 1 hour. Sections were incubated overnight at 4°C with the following primary antibodies: DDIT4L (1:500; Sigma; SAB1408239); DDIT4L (1:200; Invitrogen; PA5-48874); SOX2 (1:200; STEMCELL; 60055); Olig2 (1:200; Santa Cruz; sc-48817); GFAP (1:200; Sigma; G9269); Nestin (1:500; STEMCELL; 60091); CD133 (1:200; Chemicon; MAB4310); ATP5A (1:500; Abcam; ab14748); HIF1α (1:100; Sigma; SAB2702132) and CD31 (1:50; Sigma; SAB5500059). On the next day sections were incubated with biotinylated secondary antibodies, followed by signal amplification and visualization with the avidin–biotin complex (ABC) system and DAB substrate (Vector Laboratories). Leica SP8 microscope was used for imaging.

### Proximity Ligation Assay (PLA)

Adherent cell deposited on glass slides have reached the desired confluence and pre-treated with 4% PFA for 15 mins at room temperature. Using 1×Blocking solution to block and incubate the slides in a humidity chamber for 60 minutes at 37°C. Slides were incubated overnight at 4°C with the following primary antibodies: Myc (1:2000; Sigma; C3956); HA (1:2000; Sigma; H3663). On the next day the slides were incubated with PLUS and MINUS PLA probes (1:5; Sigma; DUO92001 and DUO92005) for 1 hour at 37°C. Ligase the slides with 1X Ligation Buffer (1:40; Sigma; DUO92008) and incubate the slides for 30 minutes at 37°C. The slides were incubated with Amplification-Polymerase solution (1:80; Sigma; DUO92008) for 100 mins at 37°C. Mount the slides with a cover slip using a minimal volume of Duolink In Situ Mounting Medium with DAPI (Sigma; DUO82040). Leica SP8 microscope was used for imaging.

### Quantification of Mitochondrial Morphology

Quantification of mitochondrial morphology followed the method reported before^15^. In brief, the quantification was scored from more than 150 cells with three replicates, and it was done by a person who was blinded to the treatment of the samples. Morphology was classified as follows: “fragmented”, the majority of mitochondria were less than 5μm; “intermediate”, the majority of mitochondria were more than 5μm; and “tubular”, highly interconnected mitochondria with fewer than 4– 5 free ends.

### Immunogold TEM

Sections (90 nm) were mounted on 100Lmesh nickel Formvar-carbon coated grids, incubated with PBS containing 0.1M Glycine (Sigma-Aldrich) and 1% Bovine Serum Albumin (Sigma-Aldrich). Sections were incubated with myc antibody over night at 4°C, washed extensively with PBS (5L×L10Lmin) and exposed to secondary antibody gold (15nm, Sigma-Aldrich) for 2 hours at RT. Lastly, specimens were stained with 2% uranyloxalicacetate (pH 7, Sigma-Aldrich) for 5 min at RT and Rinse grids extensively in H_2_O and view at 80 kV in a transmission electron microscope. In total 50 fields containing 1–3 cells were used for a quantitative analysis of the samples. The labeling ratio were determined by counting the intracellular gold particles and gold particles outside or inside the mitochondria membrane.

### Western blots, Antibodies, and siRNA Transfection

The tissue and cells were lysed in ice-cold buffer (50 mM Tris, 150 mM NaCl, 0.1% Triton-100, 10% glycerol, 0.5 mg/ml BSA and protease inhibitors). After measuring protein concentration, western blot samples were prepared for SDS-PAGE. Antibodies purchased from Santa Cruz include Tom20 (sc-11415), from Cell Signaling include cleaved Caspase 3 (#9661s), phospho-S6 (#9234s), S6 (#2708s), phosphor-Akt (#9271s) from Sigma include DDIT4L (SAB1408239 or SAB2100546), HIF1α (SAB2702132), HA-tag (H3663), myc-tag (C3956), Flag-tag (F3165), from Proteintech include myc-tag (60003-2-Ig), GAPDH (10494-1-AP), from Abcam include ATP5A (ab14748), ATP5B (ab14730), from Amersham pharmacia include GST (#27-4577). Random shuffle sequence for control and a DDIT4L siRNA (DDIT4L-homo-620 sequence: GUGUGAUUCUAGCGUCGUA) from GenePharma was used to knockdown experiments in GBM cell lines. Lipofectamine® 3000 transfection reagent (Thermofisher, L3000015) was used for siRNA transfection according to manufacturer’s instructions.

### Immunoprecipitation and GST pull-down

The GBM primary cells (hGBM11879), U87MG, COS7, HEK293 and HEK293T cells were lysed in ice-cold buffer (50 mM Tris, 150 mM NaCl, 0.1% Triton-100, 10% glycerol, 0.5 mg/ml BSA and protease inhibitors). The suspended lysate was immunoprecipitated with antibodies for 1 hour at 4°C and then with Protein G-Agarose for Rb or Mo antibodies overnight at 4°C. Immunoprecipitates were collected and aspirated. The sepharose was resuspended in RIPA buffer without SDS, washed at least 3 times, and incubated in SDS buffer for 20 min at 60°C. Then the immunoblotting was processed. The immunoblotting band in U87MG cells was analyzed for mass spectrometry. The GST-fused protein (5 mg) was incubated with the cell lysate at 4°C overnight. Then, the GST-fused protein was precipitated with 10 mL of Glutathione-Sepharose beads at 4°C for 2 hours. The precipitant was washed, denatured and prepared for immunoblotting.

### Homology Modeling and *in-silico* Docking hDDIT4L into hTOM40 and hATPase

The homology model of human DNA damage inducible transcript 4-like (hDDIT4L) dimer was created based on the crystal structure of human REDD1, a hypoxia-induced regulator of mTOR (3LQ9, 2Å)^16^ using Modeller^17^. The sequence of hDDIT4L was retrieved from the Uniport (entry: Q9NX09). According to the secondary structure information of the template, the sequence alignment was adjusted manually to obtain a more reasonable alignment. The homology model of human TOM40 was created from structure of the TOM core complex from neurospora crassa^3^. The homology model of human ATP synthase was created based on the ground state structure of F1-ATPase from bovine heart mitochondria (2JDI, 1.9Å)^18^. The constructed homology model was checked and validated by the program Procheck^19^. hDDIT4L/hTOM40 and hDDIT4L/hATPase interaction models were predicted by ZDOCK^20^. 2000 possible binding poses were obtained and clustered, and then the selected best pose was optimized by DESMOND using the default parameters^21^.

### Assay for Cellular ATP Levels

The U87 cells were seeded in 96-well plates at 5 x 10^4^ cells per well and transfected with DDIT4L and control vector for 30 hours in triplicate. ATP levels were measured using the ATPlite 1step general assay (Perkinelmer, 6016739); luminescence was read using Spectramax^@^ iD3 (Molecular Devices). Readings were normalized to cell number.

### Mitochondrial Membrane Potential Measurement

The JC-10 mitochondrial membrane potential assay kit was purchased from Abcam (112134) and the assay was performed according to the manufacturer’s protocol. Cells were transfected with DDIT4L and control vector for 30 hours, then measuring of mitochondrial membrane potential with Spectramax^@^ iD3 (Molecular Devices).

### Oxygen Consumption Rate (OCR)

OCR was measured by Seahorse Bioscience instrument (XF24/XF96, Agilent) with 80-90% confluent cells according to the manufacturer’s protocol. Briefly, on the day following cell seeding, cells were equilibrated for 1 hour in a 37°C incubator lacking CO_2_. Oxygen concentration in media was measured at basal conditions and after sequential addition of compounds and peptides as indicated in corresponding figure legends. Concentration of compounds and peptides used were: oligomycin A (1 μM), Fccp (1 μM), a mixture of rotenone (500 nM) and antimycin A (500 nM), TAT-DDIT4L^V125-P132^ or scramble peptides as described in specific figure legend. A minimum of three wells were utilized per condition to calculate OCR.

### Bioinformatics Analysis

TCGA diffuse glioma database^22^ including RNA-seq, simple somatic mutation and patient survival were downloaded from the TCGA Research Network by R Studio Version 1.2.5033 with packages TCGA biolinks^23^. Gene expression of DDIT4L and other genes in the TCGA diffuse glioma database were processed using publicly available tools including the biostatistics program R Studio Version 1.2.5033 with packages gplots, DT, dplyrand Summarized Experiment. DDIT4L expression level and clinical data of Chinese Glioma Genome Atlas were downloaded from online database (CGGA, http://www.cgga.org.cn). In addition, DDIT4L expression level of Gravendeel dataset and Rembrandt dataset were obtained from the online Visualization Tools for Glioma Datasets (GlioVis, http://gliovis.bioinfo.cnio.es).

### Statistical Analysis

The data are presented as mean ± SEM. Sample number (n) values are indicated in figures, figure legends or results section. Two groups were compared by a two-tailed, unpaired Student’s t test. Comparisons between two groups with multiple time were performed by a two-way ANOVA with Bonferroni’s post hoc test. Survival analysis was performed using the Kaplan-Meier method and comparisons were done using the log-rank test.

Statistical analysis was performed using PRISM (GraphPad Software). Differences were considered significant at p < 0.05 (*p < 0.05, **p < 0.01, ***p < 0.001).

## Results

### DDIT4L expression is associated with glioma aggressiveness

Differentially expressed genes were analyzed among different types of gliomas stratified according to their clinical features to identify potential candidate genes associated with the prognosis or treatment of GBM. First, patients with GBM have a lower survival rate than patients with low-grade glioma (LGG). Second, patients with GBM expressing wild-type IDH have a worse prognosis than patients with GBM expressing mutant IDH. Last, based on prior naming and the expression of signature genes, some functional subtypes of GBM (proneural, neural, classical, and mesenchymal) have different prognoses. The proneural type of GBM imparts a survival advantage, while the mesenchymal type of GBM has a less favorable prognosis^24-26^. We focused on the up-regulated genes from TCGA database by screening the overlapping genes in three paired classifications, namely GBM & LGG, IDH mutation & wild type GBM and mesenchymal & proneural types of GBM. We identified 53 up-regulated genes by performing a comparative analysis (Fig. 1A; Table S1). The top 10 most up-regulated genes from the GBM & LGG comparison are shown (Fig. 1B, 1C). *DDIT4L* was one of the genes showing the most significant up-regulation in the IDH wild-type and mesenchymal type of GBM (Fig. S1A), while its homolog *DDIT4* was expressed at high levels in LGG (Fig. S1B). In order to find out the expression of *DDIT4L* in different types of glioma tumor tissues, we analyzed data from two different databases and found that its expression in gliomas with better prognosis, such as oligodendroglioma and astrocytoma, was lower than that in adjacent tissues, but slightly higher in GBM (Fig. S1C). As shown in CGGA database, while patients with low grades glioma presenting high level *DDIT4L* expression exhibited shorter median survival time, but patients with grade IV glioma presenting different *DDIT4L* exhibited similar survival times (Fig. S1D). Similarly, in TCGA database, patients with GBM presenting different *DDIT4L* expression levels exhibited similar survival times (Fig. S1E). Taken into the consideration that the different effects of DDIT4L in LGG and GBM patients, there is a high possibility that highly expressed DDIT4L in GBM maybe a favorable factor for the survival time of GBM patients. In a line with the bioinformatic analysis, immunostaining results showed that the DDIT4L expression level in GBM was higher than that in tissues from patients with LGG (Fig. 1D). In the GBM tissue, DDIT4L partially co-localized with the astrocyte marker GFAP (65% of DDIT4L-positive cells) or oligodendrocyte marker Oligo2 (42% of DDIT4L-positive cells) (Fig. 1E). DDIT4L was also expressed in Nestin-, CD133- and SOX2-positive glioma-initiating cells (Fig. 1F, 1G and S1F). Therefore, DDIT4L was expressed at a much higher level in patients with GBM than in patients with other gliomas.

**Fig. 1.**
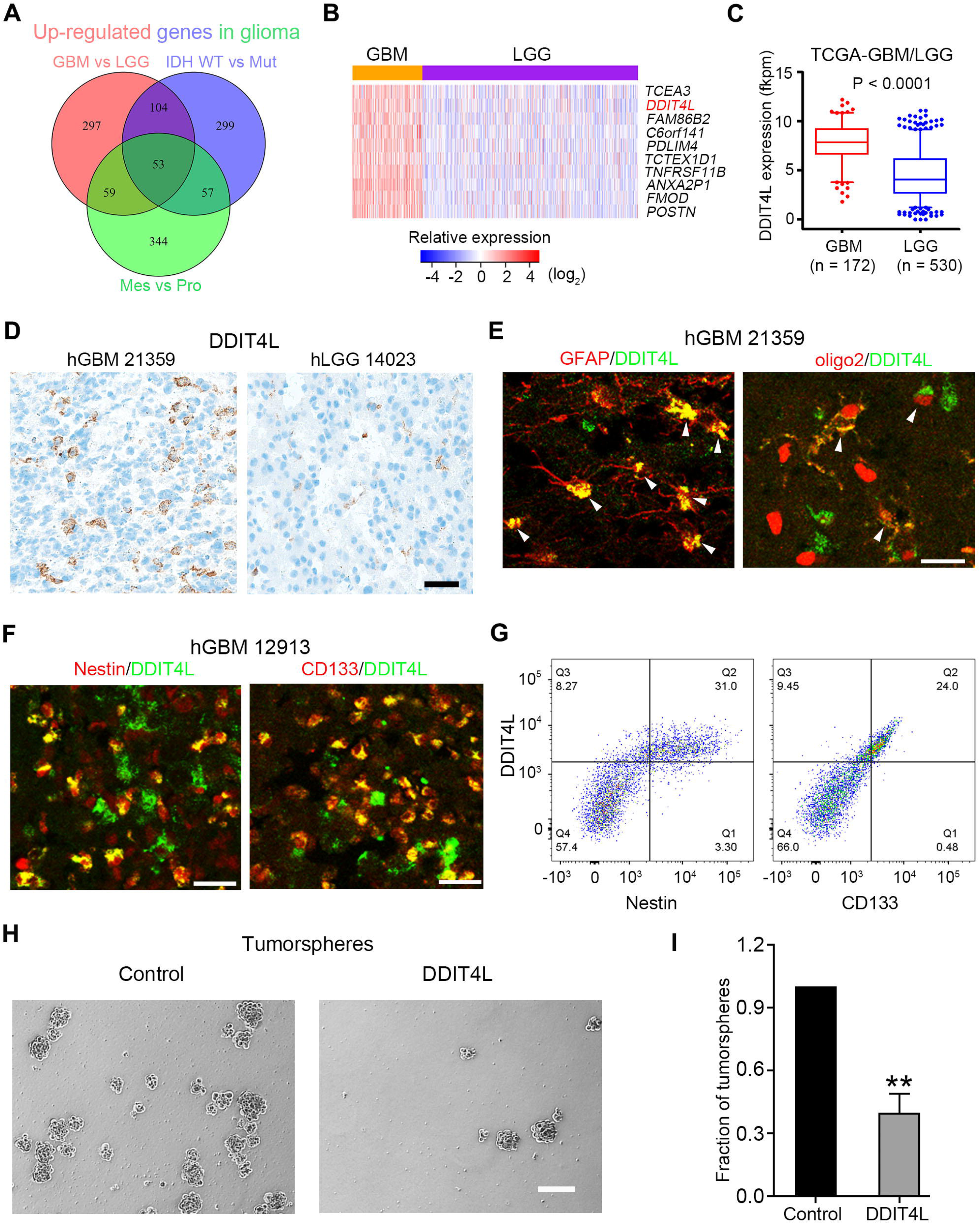
DDIT4L is expressed in aggressive glioma and suppresses tumorsphere formation. **A.** The upregulated genes from the TCGA database via overlapping GBM &LGG, IDH wild-type & mutation GBM, and mesenchymal & proneural type of GBM. **B.** Top 10 upregulated genes are identified in the GBM&LGG comparative analysis. **C.** Boxplot of the DDIT4L expression in GBM &LGG from the TCGA database. **D.** Immunohistochemical staining of DDIT4L in GBM and LGG. Representative images are shown. Scale bar = 100 μm. **E.** Double-immunostaining of DDIT4L and GFAP, DDIT4L and Oligo2 in GBM tissues. Scale bar = 50 μm. **F.** Double-immunostaining of DDIT4L and Nestin, or DDIT4L and CD133 in GBM tissues. Scale bar = 100 μm. **G.** DDIT4L partially co-expressed with Nestin and CD133 by the flow cytometry. **H, I.** Lentiviral infection of DDIT4L apparently inhibits the tumorsphere formation in the primary GBM cells. Scale bar = 200 μm.

### DDIT4L mainly localizes in hypoxia-sensitive GBM cells and suppresses tumor sphere formation

Because hypoxia is a feature of GBM and hypoxia up-regulates *DDIT4L* expression^27^, the DDIT4L protein level in patients with GBM was higher than that in patients with LGG, and the DDIT4L level in primary GBM cells increased gradually after 36 hours of hypoxic culture (Fig. S2A and S2B). In GBM tissues, DDIT4L was mainly co-expressed with hypoxia-inducible factor 1α (HIF1α, 65%) but not with CD31, a marker of endothelial cells (Fig. S2C). Flow cytometry revealed that 36.3% of all primary GBM cells were both DDIT4L-positive and HIF1α-positive, while only 5.01% of the tested primary GBM cells were both DDIT4L- and CD31-positive (Fig. S3A). Unexpectedly, DDIT4L over-expression inhibited the proliferation of U87MG cells, while RNAi targeting *DDIT4L* increased cell proliferation (Fig. S3B and S3C). Moreover, the over-expression of DDIT4L via lentiviral transfection significantly impaired the tumor sphere formation capacity of primary GBM cells (Fig. 1H and 1I). Based on these results, DDIT4L mainly localizes in hypoxia-sensitive GBM cell and suppresses GBM tumor growth.

### DDIT4L could bind to TOM40 protein

As DDIT4L inhibited GBM, except for targeted mTOR pathway, we further identified whether novel targets of DDIT4L action in the tumor cell. Such as U87MG cell line, which was used due to its similar characteristics to GBM, while DDIT4L was expressed at a low level in U87MG cells, we transfected DDIT4L tagged with myc and His, then immunoprecipitated the cell lysate with the myc antibody. Two immunoreactive bands with molecular weights of ∼40 and ∼50 kDa were coimmunoprecipitated with the myc antibody (Fig. 2A and 4A). The coimmunoprecipitated bands were purified and analyzed using mass spectrometry. This analysis showed that among the significant protein targets, 3-12 peptides matched the human TOM40 protein (∼40 kDa), and 5-16 peptides matched the human ATP5A protein (∼50 kDa). The data from three samples represented 28% and 37% coverage of the TOM40 and ATP5A protein sequences, respectively. Co-immunoprecipitation assays and proximity ligation assays (PLAs) further confirmed that TOM40 interacted with DDIT4L in GBM tissue, HEK293 cells (Fig. 2A-2C). As a control, the generally accepted import receptors TOM20, TOM22 and TOM70 did not interact with DDIT4L (Fig. 2D). Next, the Co-immunoprecipitation assays and immunostaining showed that DDIT4L interacted with and co-expressed with TOM40 in GBM cells (Fig. 2E and 2F). Then, we investigated the possible docking site of DDIT4L on TOM40 using a combination of homology modeling and *in silico* docking.

**Fig. 2.**
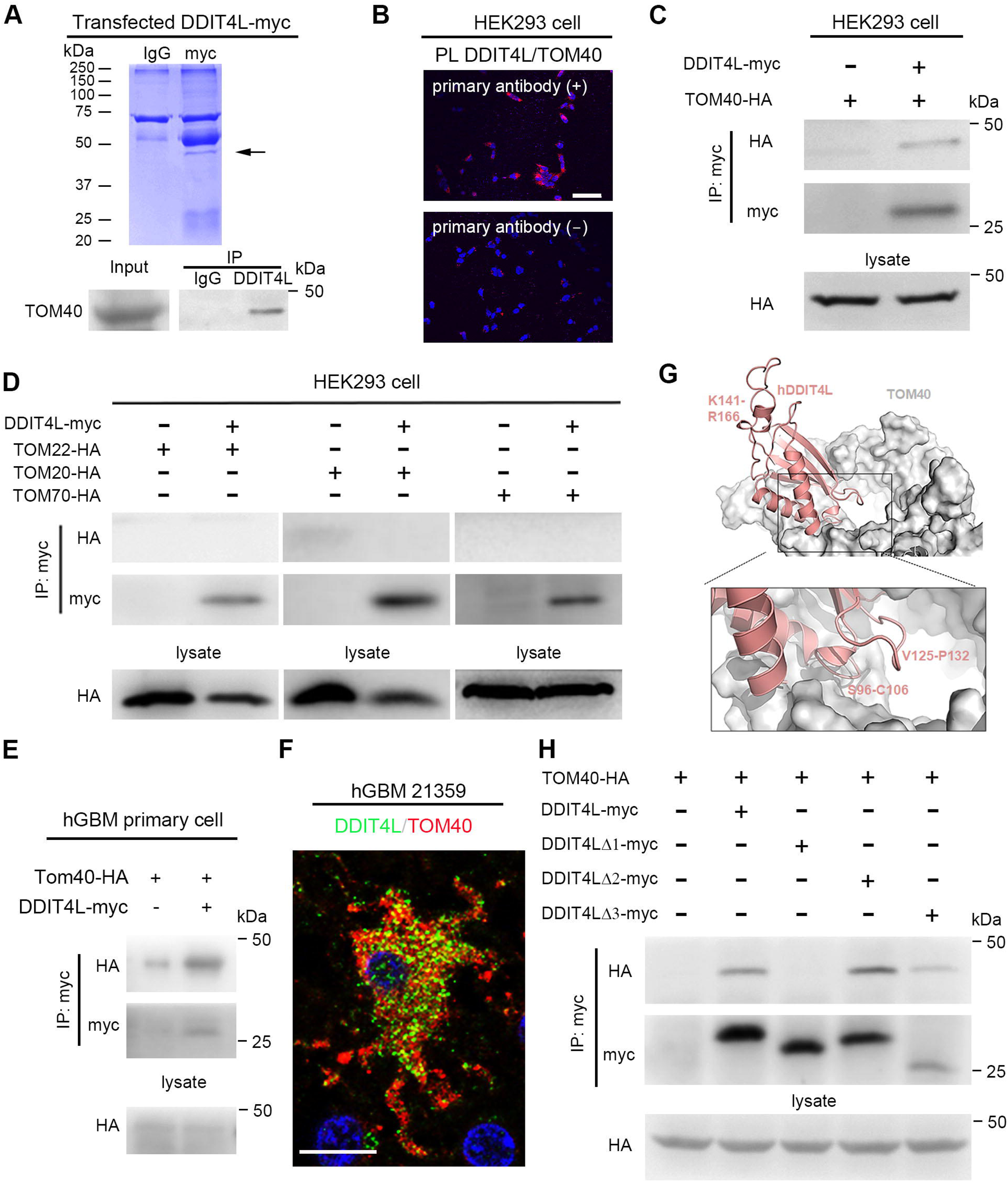
DDIT4L binds to TOM40. **A.** Coomassie blue staining shows a band of molecular weight ∼40 kDa in the myc-antibody-precipitated proteins from U87MG cells transfected the DDIT4L-myc plasmid. Co-IP shows that TOM40 (∼40 kDa) in DDIT4L-precipitated proteins from GBM tissue (n = 3). **B.** PLA assay shows that DDIT4L binds to TOM40 in HEK293 cells. Scale bar = 20 μm. **C.** In the lysate of HEK293 cells co-transfected with plasmids expressing TOM40-HA and DDIT4L-myc, TOM40-HA is found in the proteins precipitated with myc antibodies (n = 3). **D.** Co-IP shows that TOM20, TOM22 and TOM70 were not in myc-antibody precipitated proteins from HEK293 cells expressing DDIT4L-myc (n = 3 for all experiments). **E.** In the lysate of GBM primary cells (hGBM11879) co-transfected with plasmids expressing TOM40-HA and DDIT4L-myc, TOM40-HA is found in the proteins precipitated with myc antibodies (n = 3). **F.** Double immunostaining shows that DDIT4L is colocalized with TOM40 in GBM patient’s tissue. Scale bar = 20 μm. **G.** Possible docking site of DDIT4L in the TOM40 using a combination of homology modeling and *in-silicon* docking. **H.** In the lysate of HEK293 cells expressing TOM40-HA and DDIT4L-myc or DDIT4LΔ2-myc (Δ2: deleting region of V125-P132) or DDIT4LΔ3-myc (Δ3: deleting region of K141-R166), the HA signal is found in proteins precipitated with myc antibodies. The Co-IP signal of HA is decreased in the cell coexpressing TOM40-HA and DDIT4LΔ1-myc (Δ1: deleting region of S96-C106) (n = 3).

In this model, we screened 3 possible motifs, S96-C106, V125-P132 and K141-R166, in DDIT4L as possible binding sites for TOM40 (Fig. S4). The structural analysis using *in silico* docking showed that the S96-C106 domain of DDIT4L was a functional domain binding to TOM40 (Fig. 2G). Consistently, a co-immunoprecipitation assay showed that the S96-C106 motif, but not the V125-P132 and K141-R166 motifs, in DDIT4L bound to TOM40 (Fig. 2H).

### DDIT4L is transported into mitochondria via a single TOM40 protein

Next, we examined the subcellular distribution of DDIT4L in HEK293 cells. Immunostaining results showed that DDIT4L did not co-localize with mitochondria labeled with a mitochondrial tracker (MitoTracker) in HEK293 cells transfected with DDIT4L (Fig. S5A). Surprisingly, in HEK293 cells co-transfected with DDIT4L and TOM40, DDIT4L mainly co-localized with mitochondria (Fig. S5B and S5C). Furthermore, DDIT4L did not localize to the inner mitochondrial membrane labelled by ATP5B 24 h after transfection in COS7 cells transfected with DDIT4L. Consistently, in COS7 cells co-transfected with DDIT4L and TOM40, DDIT4L mainly localized in TOM40-positive granules 12 h after transfection, and the localization in the ATP5B-positive inner mitochondrial membrane appeared (Fig. 3A and 3B). The majority of DDIT4L-positive puncta were localized in TOM40-positive mitochondria 24 h after transfection (Fig. 3A and 3B). Moreover, DDIT4L mainly co-localized with the lysosome marker LAMP1 in HEK293 cells transfected with DDIT4L. In HEK293 cells co-transfected with DDIT4L and TOM40, DDIT4L mainly co-localized with MitoTracker (Fig. 3C and 3D). Thus, DDIT4L was transported into mitochondria by interacting with a single TOM40 molecule, not the TOM complex. Immunoelectron microscopy showed that in HEK293 cells transfected with DDIT4L, immunogold labeling of DDIT4L was mainly dispersed in the cytoplasm, and the morphology of mitochondria seemed normal. In HEK293 cells co-transfected with DDIT4L and TOM40, immunogold labeling of DDIT4L mainly appeared in mitochondria, and the mitochondria swelled and showed a reduced number of cristae (Fig. 3E and 3F). Taken together, these results indicated that a single TOM40 protein on the outer membrane of mitochondria interacted with and transported DDIT4L into the mitochondria, resulting in the disfunction of mitochondria.

**Fig. 3.**
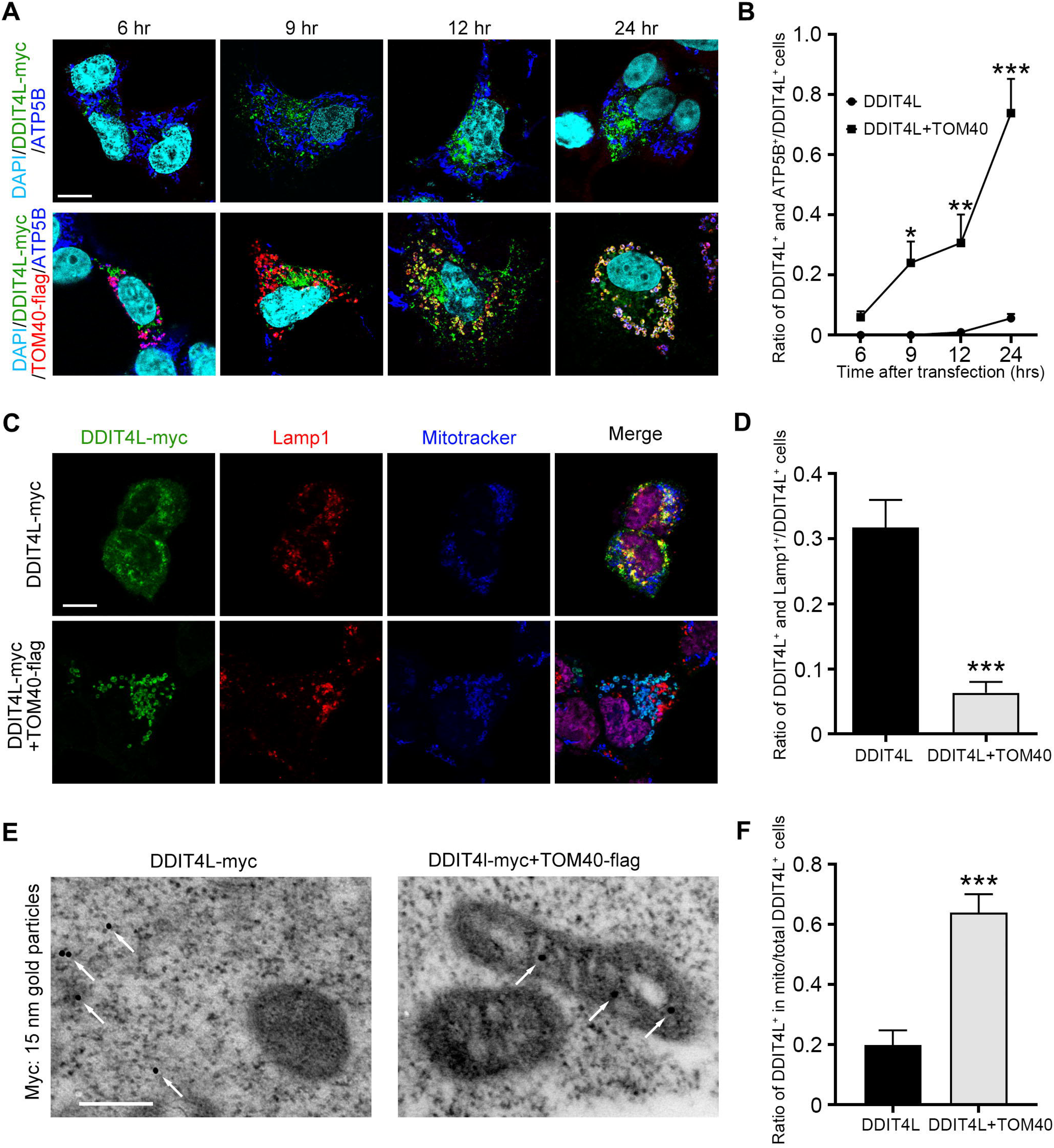
DDIT4L transports into mitochondria via TOM40. **A.** Transfected DDIT4L did not localize in mitochondria after 24h, co-transfected DDIT4L and TOM40, DDIT4L mainly localized in TOM40-postive granule at 12h, and the ATP5B positive inner membrane of mitochondria have disappeared. Scale bar = 10 μm. **B.** Quantification shows that the ratio of DDIT4L positive and ATP5B positive cells/DDIT4L positive cells. **C.** Transfected DDIT4L co-localized with LAMP1, not co-localize with Mitotracker, while co-transfected DDIT4L and TOM40, the DDIT4L mainly co-localized with Mitotracker, not co-localize with LAMP1 in the HEK293T cells. Scale bar = 10 μm. **D.** Quantification shows that the ratio of DDIT4L positive and Lamp1 positive cells/DDIT4L positive cells. **E.** Electron microscopy showed that transfected DDIT4L in the HEK293 cells mainly dispersed in cytoplasm. While after co-transfection of DDIT4L and TOM40, DDIT4L mainly appeared in mitochondria and morphology of mitochondria showed swelling and a reduction in crista. Scale bar = 200 nm. **F.** Quantification shows that the ratio of DDIT4L immunogold granule in mitochondria/total DDIT4L immunogold granule in cell body.

### DDIT4L interacts with ATP5A

DDIT4L is transported via TOM40 into the inner mitochondrial membrane of mitochondria, where DDIT4L could also interact with the ATP5A according to the mass spectrometry (Fig. 2A, 4A). A co-immunoprecipitation assay showed that ATP5A interacted with DDIT4L-myc (Fig. 4A). Immunostaining showed that DDIT4L was coexpressed with ATP5A in GBM cells (Fig. 4B). Furthermore, we co-transfected plasmids expressing DDIT4L with ATP5A or ATP5B in GBM primary cells, co-immunoprecipitation assay actually showed that both ATP5A and ATP5B could interact with DDIT4L *in vitro* (Fig. 4C), which maybe induced by endogenous ATP5A or ATP5B interacting with exogenous ATP5B or ATP5A. The MS and GST pull-down experiments confirmed DDIT4L interacted with ATP5A (Fig. 4A and 4D), and purified protein interaction indicated ATP5A is a direct target of DDIT4L (Fig. 4E). We next investigated the key motifs that were involved in the DDIT4L-mediated modulation of ATPase activation to determine the precise region binding to ATP5A. We first identified a possible docking site of DDIT4L in the ATPase using a combination of homology modeling and *in silico* docking. In this model, we screened the same S96-C106, V125-P132 and K141-R166 motifs in DDIT4L as possible binding sites for ATP5A (Fig. S6A). The V125-P132 and K141-R166 motifs in DDIT4L but not the S96-C106, were the potential major binding sites for ATP5A (Fig. 4F). We also predicted that five motifs (V118-V131, K132-I158, I179-G192, L471-A486, and G491-I504) of ATP5A might also be binding domains for DDIT4L (Fig. S6B). We constructed a series of truncated ATP5A proteins fused with HA tags. Using a GST pull-down assay, the K132-I158 motif in ATP5A was identified as the major binding domain for DDIT4L (Fig. 4G). Furthermore, the interaction between DDIT4L and ATP5A was disturbed by the synthesized TAT-ATP5A^K132-I158^ peptide (Fig. S6C). The structural analysis using *in silico* docking showed that the V125-P132 domain of DDIT4L was a functional domain binding to ATP5A (Fig. 4H). In summary, DDIT4L and ATP5A could interact each other via the specific motifs.

**Fig. 4.**
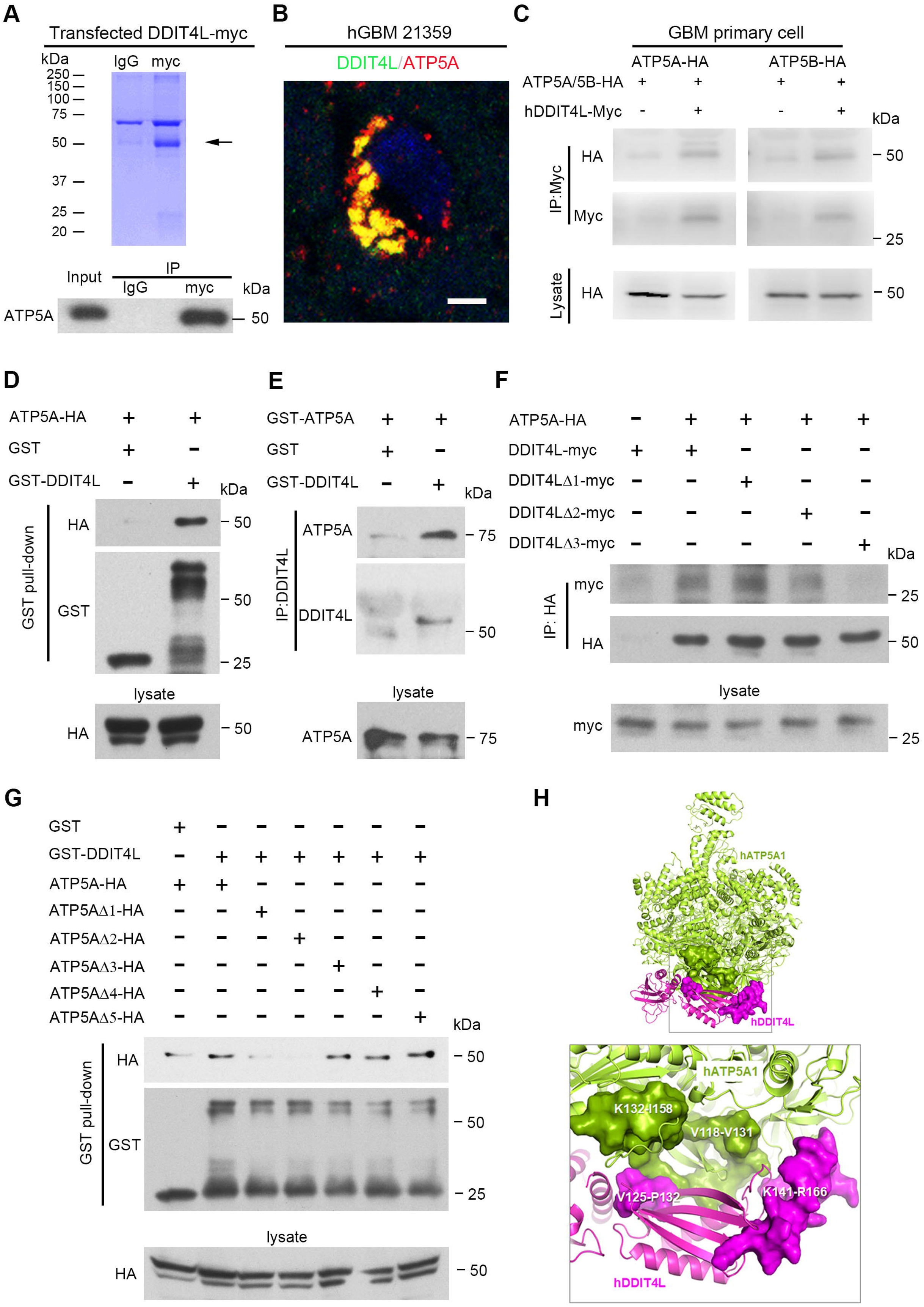
DDIT4L interacts with ATP5A. **A.** Coomassie blue staining shows a band of molecular weight ∼50 kDa in the myc-antibody-precipitated proteins from U87MG cells transfected the DDIT4L-myc plasmid. Co-IP shows that ATP5A (∼50 kDa) in myc-antibody-precipitated proteins from U87MG cells expressing DDIT4L-myc (n = 3). **B.** Double immunostaining shows that DDIT4L is co-localized with ATP5A in GBM patient’s tissue. Scale bar = 20 μm. **C.** In the lysate of GBM primary cells (hGBM11879) co-transfected with plasmids expressing ATP5A-HA or ATP5B-HA and DDIT4L-myc, both ATP5A-HA and ATP5B-HA are found in the proteins precipitated with myc antibodies (n = 3). **D.** The GST pull-down assay shows that GST-DDIT4L protein interacts with ATP5A-HA (n = 3). **E.** Co-IP with two purified proteins, GST-ATP5A and GST-DDIT4L, shows the direct interaction between ATP5A and DDIT4L (n = 3). **F.** In the lysate of HEK293T cells expressing ATP5A-HA and DDIT4L-myc or DDIT4LΔ1-myc (Δ1: deleting region of S96-C106), the myc signal is found in proteins precipitated with HA antibodies. The Co-IP signal of myc is decreased in the cell co-expressing ATP5A-HA and DDIT4LΔ2-myc (Δ2: deleting region of V125-P132) or DDIT4LΔ3-myc (Δ3: deleting region of K141-R166) (n = 3). **G.** The GST pull-down assay shows that GST-DDIT4L protein interacts with ATP5A-HA, ATP5AΔ3-HA, ATP5AΔ4-HA and ATP5AΔ5-HA, but the interaction is decreased in ATP5AΔ1-HA and ATP5AΔ2-HA in HEK293T cells (Δ1: deleting region of V118-V131, Δ2: deleting region of K132-I158, Δ3: deleting region of I179-G192, Δ4: deleting region of L471-A486, Δ5: deleting region of G491-I504) (n = 3). **H.** The selected DDIT4L (purple) /ATP synthase (green) model is derived from clustering analysis of 2000 possible binding poses based on the ZDOCK scoring, and following optimization using molecular dynamics simulations. Possible motifs implicated in interactions between DDIT4L and ATP synthase are highlighted using purple and green surface modes, respectively.

### DDIT4L inhibits mitochondrial function and induces cell apoptosis

Next, we examined the role of DDIT4L in mitochondrial function. A greater increase in *TOMM40* mRNA levels was observed than the increase in *TOMM22* or *TOMM20* mRNA levels in GBM (Fig. S7A). CGGA dataset demonstrated that *TOMM40* expression level increased in WHO grade III and IV when compared to that in Grade II gliomas (Fig. S7B). Therefore, DDIT4L should have a chance to interact with the endogenous single TOM40 protein in GBM cells. Then, DDIT4L over-expression decreased the cellular ATP content and mitochondrial membrane potential in U87MG cells (Fig. 5A and 5B). As expected, DDIT4L reduced both the basal and ATP synthase-linked oxygen consumption rates (OCRs) (Fig. 5C) and induced mitochondrial fusion, as shown by staining with a mitochondrial tracker (MitoTracker), in primary GBM cells (Fig. 5D). Electron microscopy in primary GBM cells showed an increase in mitochondrial fusion and a reduction in the number of cristae (Fig. 5E).

**Fig. 5.**
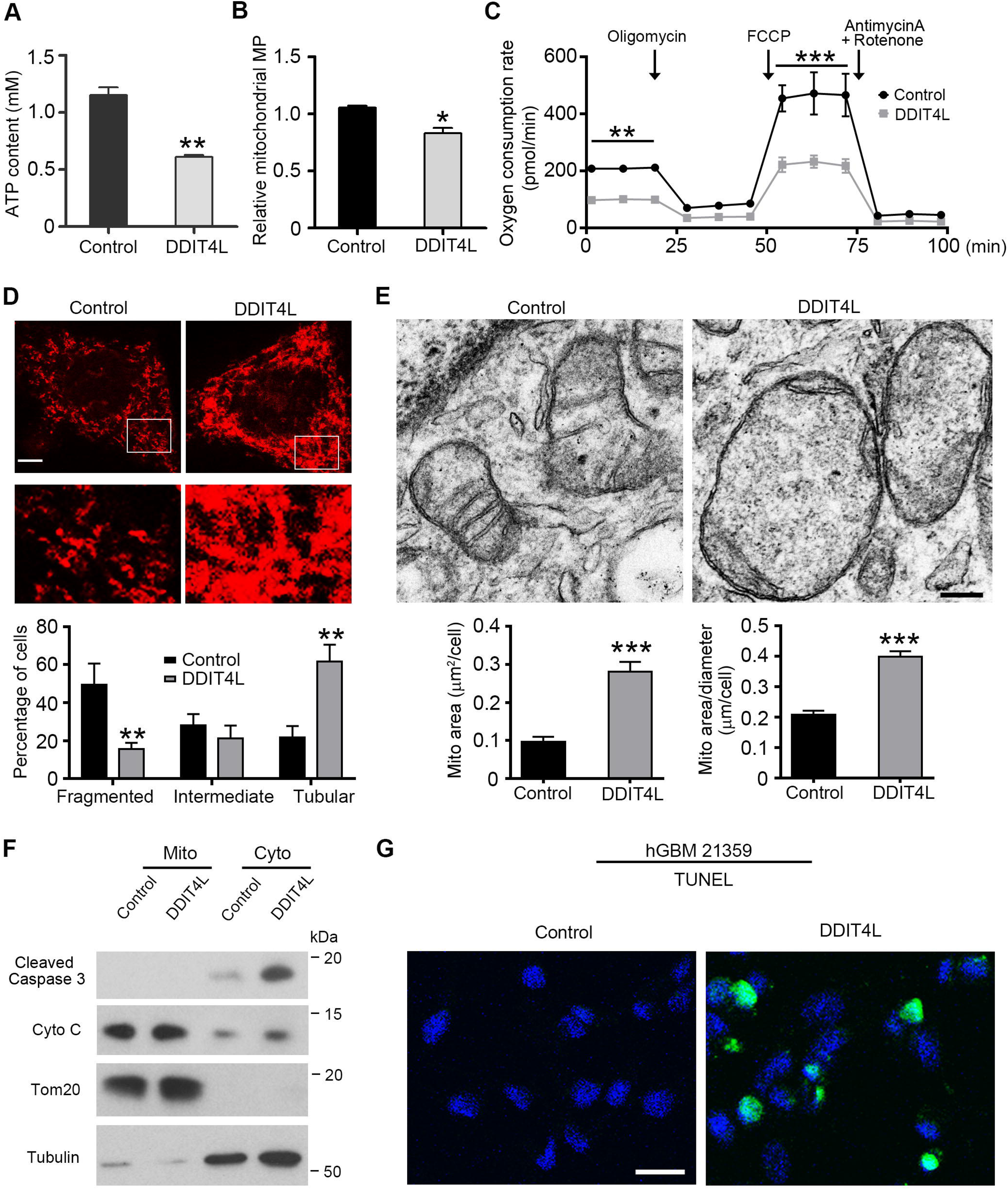
DDIT4L suppresses ATP synthase, and induces cell apoptosis. **A.** Decreased cellular ATP content in tumor cells transfected with the plasmid expressing DDIT4L. **B.** Decreased mitochondrial membrane potential in tumor cells transfected with the DDIT4L plasmid. **C.** Seahorse Mito Stress Kit assay showed that oxygen consumption rates (OCRs) on tumor cells transfected with the DDIT4L plasmid decreased when compared with that transfected with control plasmid (n = 3). **D.** Mitochondrial morphology is shown by a mitochondrial marker Mitotracker in the GBM cells transfected with DDIT4L plasmid. Scale bar = 10 μm. DDIT4L apparently increases the tubular-like mitochondria in cells. **E.** Representative EM images of the GBM cells transfected with the plasmid expressing DDIT4L show the mitochondrion fusion and reduction of mitochondrion crista after expressing DDIT4L. Scale bar = 200 nm. Quantification shows that the expressing of DDIT4L increases the mitochondria area and the ratio of area/diameter. **F.** The protein level of cytochrome c and cleaved caspase 3 in the cytoplasm are increased via isolating mitochondria from the cytoplasm of GBM cells transfected with the plasmid expressing DDIT4L (n = 3). **G.** The TUNEL assay shows that DDIT4L induces cell apoptosis (green) in primary GBM cells (DAPI, blue). Scale bar = 50 μm.

DDIT4L decreased the levels of cytochrome c in mitochondria in primary GBM and U87MG cells, while cytochrome c and cleaved caspase 3 levels were increased in the cytoplasm (Fig. 5F and S7C). DDIT4L over-expression in differentiated human astrocytes and COS7 cells did not change the levels of cytochrome c and cleaved caspase 3 in the mitochondria and cytoplasm (data not shown). Thus, DDIT4L specifically induced apoptosis in primary GBM cells, as shown by the TUNEL assay (Fig. 5G).

### Over-expressed DDIT4L and the TAT-DDIT4L^V125-P132^ peptide suppress oncogenesis of GBM

In the patient-derived and U87MG cell orthotopic mouse model of GBM, DDIT4L over-expression induced by a lentivirus reduced the tumor size and extended the lifespan (Fig. 6A-6C and Fig. S8A-S8C). Because the V125-P132 and K141-R166 domains in DDIT4L were major regions binding to ATP5A, we next examined the effects of TAT-DDIT4L^V125-P132^ and TAT-DDIT4L^K141-R166^ peptides on cell proliferation. The TAT-DDIT4L^V125-P132^ peptide, but not TAT-DDIT4L^K141-R166^ peptide, reduced the proliferation of tumor cells in a dose-dependent manner (Fig. 6D). The TAT-DDIT4L^V125-P132^ peptide also reduced the ATP synthase-linked oxygen consumption rates (OCRs) in tumor cells (Fig. 6E).

**Fig. 6.**
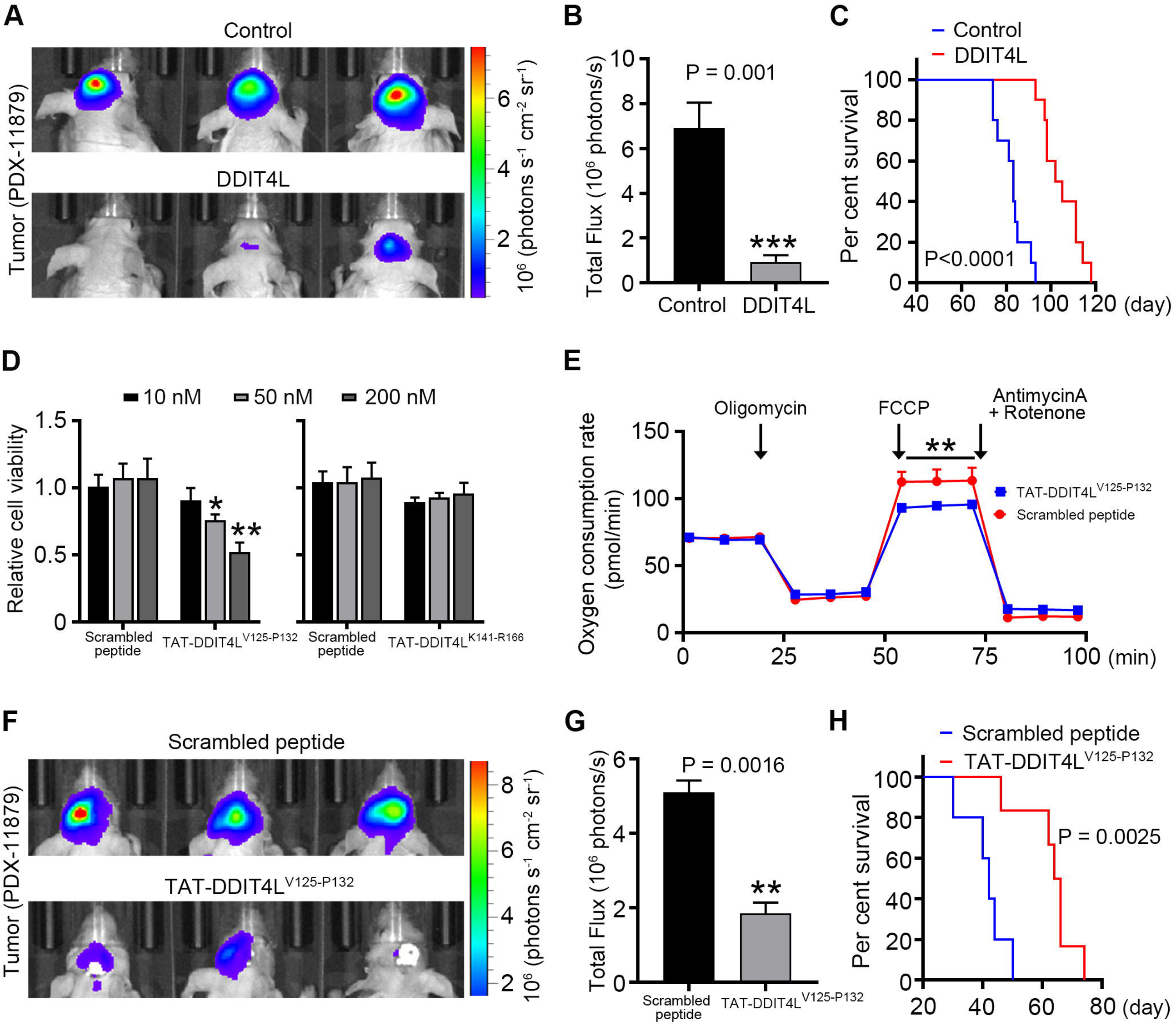
DDIT4L and TAT-DDIT4L^V125-P132^ suppress oncogenesis of GBM. **A.** Bioluminescence imaging of GBM xenografts derived from patient tumor cells expressing the luciferase and the lentivirus vector control or lentivirus DDIT4L show that DDIT4L inhibit tumor growth. Representative images are captured from animals on day 70 after cells intracranial transplantation. **B.** Total Flux of photon emission is decreased by DDIT4L (n = 10 per group). **C.** Kaplan-Meier survival analysis of mice with GBM xenografts after the transfection with lentivirus expressing DDIT4L shows that overexpressing of DDIT4L prolong the survival time significantly (n = 10 per group). **D.** TAT-DDIT4L^V125-P132^, but not TAT-DDIT4L^K144-R166^, produces an inhibitory effect on the U87MG cell proliferation in a dose-dependent manner (n = 3). **E.** Seahorse Mito Stress Kit assay on U87MG cells showed that TAT-DDIT4L^V125-P132^-induced OCRs decreased when compared with the scrambled peptide-induced OCRs (n = 3). **F.** Bioluminescence imaging of GBM patient derived tumor expressing the luciferase after treatment with TAT-DDIT4L^V125-P132^ or scrambled peptide show that TAT-DDIT4L ^V125-P132^ suppress tumor growth. Representative images are captured from animals on day 30 after minipumps implanted. **G.** Total Flux of photon emission is decreased by TAT-DDIT4L^V125-P132^ (n = 6 per group). **H.** Kaplan-Meier survival analysis of mice with GBM xenografts after treatment with TAT-DDIT4L^V125-P132^ or scrambled peptide shows that TAT-DDIT4L^V125-P132^ extend lifespan dramatically (n = 6 per group).

Importantly, four weeks after the primary cells were orthotopically transplanted in a patient-derived orthotopic mouse model of GBM, catheters were implanted at the injection site to deliver the TAT-DDIT4L^V125-P132^ peptide locally through the subcutaneous minipumps. Treatment for two weeks with the TAT-DDIT4L^V125-P132^ peptide reduced the tumor size by ∼65% and significantly prolonged the lifespan (Fig. 6F-6H). In conclusion, in both patient-derived and U87MG cell orthotopic mouse models of GBM, exogenous excessive DDIT4L and the TAT-DDIT4L^V125-P132^ peptide inhibited GBM tumor growth and prolonged the animals’ lifespan.

## DISCUSSION

According to the WHO classification, the gliomas have different types based on the distinct criteria^28,29^. Many studies have demonstrated the diverse roles of the differentially expressed genes, DNA methylation signatures, as well as specific proteins in the gliomas^30-33^. GBM is the primary brain tumor with the highest mortality rate due to hypoxia^34^. The present study shows increased expression of DDIT4L in patients with GBM, consistent with the hypothesis that hypoxia increases *DDIT4L* expression^27^. Interestingly, our results show that DDIT4L binds TOM40 for transport into mitochondria, interacts with ATP5A at the inner mitochondrial membrane, and then inhibits ATP synthase activity. Critically, treatment with exogenous excessive DDIT4L or the DDIT4L^V125-P132^ peptide causes GBM cell apoptosis and tumor suppression, suggesting that the DDIT4L and DDIT4L^V125-P132^ peptide may have the potential for GBM therapy.

DDIT4L and its homolog DDIT4 have been identified as inhibitors of mTOR signaling^4,5,7,35^. Previous studies have shown that DDIT4 expression is increased in tumors characterized by a worse prognosis, such as acute myeloid leukemia, breast cancer, GBM, colon, skin and lung cancer^36^. The potential molecular mechanism of DDIT4 in oncogenesis is that DDIT4 binds 14-3-3 proteins to elicit tuberous sclerosis complex 2 (TSC2)/14-3-3 dissociation, and then TSC2 inhibits mTORC1, which suppresses the proliferation of tumor cells^37,38^. The role of DDIT4L in tumor growth is largely unknown, while DDIT4L suppresses mTORC1 signaling through a similar molecular mechanism in some pathological conditions, and promote autophagy^5,6^.

On the other hand, TOM40, along with TOM22 and TOM5 constitute a TOM complex that transports cytoplasmic proteins into mitochondria. Proteins first bind membrane receptors, such as TOM22, and then are transported into mitochondria via TOM40. None of the proteins directly bind TOM40 for transport into mitochondria in normal condition. Under pathological condition, TOM40 mRNA expression is apparently increased in GBM tissues, while TOM22 and TOM5 mRNA expression show no distinct changes. TOM20 mRNA expression was significantly decreased in GBM tissues, and thus, TOM40 existed as a single molecule except when it was present in the TOM complex, which provided DDIT4L an opportunity to bind TOM40. In present study, we firstly identified that DDIT4L interacted with import channel TOM40 other than import receptors in tumor cells but not in normal cells. Then, DDIT4L inhibited the activity of ATP synthase to release cytochrome c from mitochondria into the cytosol and activated the caspase pathway to elicit cell apoptosis. The comprehensive transcriptome analysis indicated high level of DDIT4L in LGG have a poor prognosis, while high level of DDIT4L in GBM have no change for prognosis compared with low level. This should be attributed to different expressions of TOM40 in LGG and GBM. For DDIT4L is a hypoxic-sensitive molecule, and our experimental findings demonstrate a robust upregulation of DDIT4L in response to hypoxic conditions. According to the bioinformatics analysis and our experimental results, LGG have a relatively high level of DDIT4L, indicating tumor tissues have apparent hypoxic condition. While in LGG, the TOM40 mostly expressed as low level as normal tissues, so DDIT4L could not bind to single TOM40 enough, and not inhibit mitochondrial function. In result, DDIT4L turns to suppress mTORC1 signaling and promote autophagy, which promotes tumor cell survival, lastly leads to poor prognosis. While GBM is a high-grade malignancies characteristic of apparent proliferation, which results to localized hypoxia^39^. This hypoxic microenvironment, in turn, prompts the expression of DDIT4L in tumor cells, and TOM40 also significantly increases expression in GBM tissues, which triggers DDIT4L binding to single TOM40, then translocating the inner membrane of mitochondria, interacting with ATP5A, lastly inhibits mitochondrial function and leads to cell death. While DDIT4L at same time also suppresses mTORC1 signaling and promote autophagy, which promotes tumor cell survival. Consideration the two conditions, endogenous DDIT4L elevation have no change prognosis in GBM, in which its microenviroment likes a balanced seesaw between the TOM40-ATP5A related mitochondrial disfunction and mTORC1 signaling. More importantly, we identified a peptide DDIT4L^V125-P132^, capable of mimicking the inhibitory effect of DDIT4L on GBM growth, in which the microenvironment seesaw falls to the side of mitochondrial disfunction, showing the potential to be developed as a therapeutic biologic for GBM treatment.

This study shows a novel molecular mechanism by which DDIT4L, not DDIT4, binds to TOM40 for transport into mitochondria, interacts with ATP5A to suppress mitochondrial ATP synthase activity, and elicits cell apoptosis. This mechanism may provide a possible new therapy for GBM. Taken together, *in vitro*, bioinformatics and *in vivo* findings identify DDIT4L as a novel endogenous suppressor of GBM by interacting with TOM40 and ATP5A. These findings provide novel insights into the relationship between mitochondrial function and cell apoptosis during GBM oncogenesis.

## Authorship

LKC and MY conceived the project. ZX, MY, CL and LKC designed experiment. YP, TQS, YBJ and QZX (primary GBM cell culture and PDX model), CB, LH, PJ and ZXY (plasmid construction, cell line culture and IP), HXY and GPP (immunostaining, ATP1step and OCR), LRJ (MS), YY (Homology Modeling and *in-silico* Docking), ZZN (hypoxia culture) completed experiments. LKC, LH and YP performed statistical analyses and made figures. YP performed informatics analyses. LKC, ZX, MY, CL, YP and LH prepared the manuscript. LKC, ZX, MY and CL revised the manuscript.

## Acknowledgements

We thank Jingyi Hui, Juyi Wen and Qingjun Meng for discussions, Hao Feng (National Facility for Protein Science in Shanghai) for animal management.

