## Supplemental information for "The DDIT4L-TOM40-ATP5A pathway suppresses glioblastoma oncogenesis"

**SUPPLEMENTARY INFORMATION**

**Cell Growth and Viability Assays**

Cells were seeded in 6-well plates, transfected with DDIT4L-myc and control plasmids or DDIT4L and control RNAi or treated with TAT-DDIT4L^V125-P132^ peptide or scrambling peptide, then harvesting cells at 0, 1, 2, 3 days, staining with trypan blue stain (0.4%, Thermofisher, T10282). Cell number and viability were measured by automated cell counter Cellometer^®^Auto 1000 (Nexcelom Bioscience) following the manufacturer’s instructions.

**Neurosphere assay**

For the sphere assay, 5000 dissociated primary cells were seeded into one well of a 6 well Ultra-Low attachment plate (Life Sciences Costar®, 3471) and grown in NSC medium for 2 weeks and imagined every other day. A picture of the whole well was taken with a Zeiss widefield microscope and the neuroshperes were counted and analyzed with ImageJ. The assay was repeated at least three times for all cell lines and the significance was analyzed with a One-Way ANOVA test.

**Flow Cytometry**

Flow cytometry was performed on a BD LSRFortessa™ X-20 cell analyzer (BD Biosciences) and analyzed using FlowJo software (Tree Star). The primary cells were incubated for 10 min at 37°C in 10 ml of DPBS (Thermofisher, 14190144) containing 0.25% trypsin (Thermofisher, 25300). The suspension was centrifuged at 800g for 5 min at 4°C. The pellet was re-suspended in DPBS and centrifuged again at 800g for 5 min at 4°C. The pellet was re-suspended in 1 ml paraformaldehyde (4%, pH=7.4) and stayed for 10 min at 4°C. The suspension was centrifuged at 800g for 5 min at 4°C. The pellet was re-suspended in DPBS and centrifuged again at 800g for 5 min at 4°C. The pellet was re-suspended in 0.1% Triton X-100 for 10 min at 4°C. The suspension was centrifuged at 1200g for 10 min at 4°C. The pellet was re-suspended in PBA (PBS consisting of 1% BSA, 0.1% Sodium azide) containing primary antibodies incubated for 1 h at 4°C. The suspension was centrifuged at 1200g for 10 min at 4°C. The pellet was re-suspended in PBA containing secondary antibodies incubated for 30 min at 4°C. The suspension was ready for testing, fluorescence-minus-one control based gating.

**Plasmid Construction**

All primers and oligonucleotides used for the construction of plasmids expressing DDIT4L and DDIT4L truncations, ATP5A and ATP5A truncations, ATP5B, GST-DDIT4L, GST-ATP5A are listed in Table S2.

The cDNA of human DDIT4L was cloned into pcDNA3.1/myc-his(+)A vector for the expression of DDIT4L-myc. The plasmids expressing DDIT4LΔ1-myc, DDIT4LΔ2-myc and DDIT4LΔ3-myc were modified from the DDIT4L-myc plasmid using KOD-plus Mutagenesis Kit (Toyobo). The cDNA of human TOM20, TOM22, TOM40, ATP5A and ATP5B was constructed in a modulated pcDNA3.1/HA-his(+)A vector as a HA tag replaced the myc tag in the pcDNA3.1/myc-his(+)A vector. The plasmids expressing ATP5AΔ1-HA, ATP5AΔ2-HA, ATP5AΔ3-HA, ATP5AΔ4-HA and ATP5AΔ5-HA were modified from the ATP5A-HA plasmid using KOD-plus Mutagenesis Kit (Toyobo). The cDNA of human DDIT4L and human ATP5A following a HA tag was cloned into a pGEX vector for the construction of the GST-DDIT4L and GST-ATP5A-HA plasmids.

**Preparation of the GST-Fused Proteins**

The GST-fused proteins were expressed in Escherichia coli BL21. The bacteria were grown in 23YTA media and the protein expression was induced by 1 mM Isopropyl b-D-thiogalactopyranoside (IPTG). Then the bacteria were centrifuged, resuspended and sonicated for the protein release. The proteins were purified using Glutathione-Sepharose beads (GE Healthcare), concentrated and quantified before use.

**Transmission electron microscope (TEM)**

The primary cells (5×10^5^) were collected after transfected with 3 μg DDIT4L or control plasmid for 48h. Then, the cells were first fixed in 2.5% Glutaraldehyde buffered in 0.1M PBS for 2h. The samples were post-fixed with 1% OsO4 in the dark for 1.5h. The samples were dehydrated through alcohol with gradient concentrations (30, 50, 70, 80, 95, and 100%) and further acetone dehydration twice (100%). Finally, the samples were treated with acetone: Epon812 (1:1) for 2h, and treated with Epon812 (100%) overnight. Then, the ultra-thin sections (70 nm) were obtained, stained with Uranium acetate and Lead citrate, and photographed under a TEM at 80kV (FEI Tecnai G2 Spirit). Mitochondria were manually segmented, the area and diameter measured in Fiji software by adding up the voxel.

**Mass Spectrometry (MS)**

For MS analysis, the in-gel digestion was performed using the following protocol. To reduce disulfide bonds, the strip was performed in 1% 1, 4-dithiotreitol (DTT) in SDS equilibration buffer [50 mM Tris-Cl (pH 8.8), 6 M urea, 30% glycerol, 2% SDS and bromophenol blue] for 15 min. This step was followed by alkylation of the free sulhydryl groups by 2.5% iodoacetamide in a SDS equilibration buffer for another 15 min in the dark at room temperature. Then the strip was cut into 18 gel sections (each section about 1.0 cm in length). Each gel section was washed three times alternately with acetonitrile and 100 mM ammonium bicarbonate. During the last wash the gel slices were incubated in 100 mM ammonium bicarbonate for 15 min at temperature 4°C. The gel slices were dried by vacuum centrifugation and allowed to swell in a 50 μl trypsin solution containing trypsin (20 μg/ml) and ammonium bicarbonate (50 mM) at 4°C for 45 min. After adding another 50 μl of trypsin solution, the gel slices were kept at 37°C for 20 h. The supernatant was transferred to another vial, and the gel slices were extracted for 15 min three times by 0.1% formic acid in 60% acetonitrile. The recovered peptide solutions were dried by vacuum centrifugation and desalted and cleaned using a Ziptip (Millipore, Corp., Bedford, MA).

The peptide mixtures from each section of the strip were separated by Reverse phase HPLC (RP-HPLC) followed by tandem mass analysis. RP-HPLC was performed on a surveyor LC system (Thermo Finnigan, San Jose, CA). The C18 column (RP, 180 μm × 150 mm) was obtained from Column Technology Inc. (Fremeont, CA,). The pump flow was split 1:120 to achieve a column flow rate of 1.5 μl/min. Mobile phase A was 0.1% formic acid in water, and mobile phase B was 0.1% formic acid in acetonitrile. The tryptic peptide mixtures were eluted using a gradient of 2−98% B over 180 min.

The MS was performed on a LTQ linear ion trap mass spectrometer (Thermo Finnigan, San Jose, CA) equipped with an electrospray interface and operated in positive ion mode. The capillary temperature was set to 170°C and the spray voltage was at 3.4 kV. Normalized collision energy was at 35%. Automatic gain control was used to obtain maximal signal of each scan. The mass spectrometer was set so that one full MS scan was followed by ten MS/MS scans on the 10 most intense ions. Dynamic Exclusion was set at repeat count 2, repeat duration 30 s, and exclusion duration 90 s.

The acquired MS/MS spectra were searched against the IPI rat database using BioWorks 3.0 software (Thermofinnigan) on an 8 node Dell PowerEdge 2650 cluster. An accepted SEQUEST result had a ΔCn score of at least 0.1 (regardless of charge state), a value known for high confidence in a SEQUEST search. All output results were combined together using homemade software named Build Summary to delete the redundant data. To make sure that the MS/MS spectrum was of good quality with fragment ions clearly above baseline noise, we referred to the parameters reported in previous studies and applied stricter criteria for peptide identification. Peptides were validated after meeting the following criteria. The SEQUEST cross-correlation score must be ≥1.9 for a +1 tryptic peptide, ≥2.2 for a +2 tryptic peptide and ≥3.75 for a +3 tryptic peptide. In addition, ΔCn cutoff values were ≥0.1 and the SP rank of the peptides ≤4.

**Magnetic resonance imaging (MRI)**

All in vivo MRI measurements were performed on a 7.0 Tesla small animal MR system (Biospec, Bruker Biospin). Mice were anesthetized by inhalation narcosis using 0.5% to 1.5% isoflurane (RWD), pressurized oxygen. Core body temperature was maintained at 37°C. Respiration rate and temperature were monitored using a remote monitoring system (Model RM400S, RWD Inc.). T2-weighted images were acquired with the same slice geometry (slice thickness = 500 µm, 25 coronal slices covering a brain region of 12.5 mm (approx. Bregma 4 to -8.5mm)). Tumor regions were manually segmented and the volume calculated in Fiji software by adding up the voxel volumes.

**FIGURES and LEGENDS**

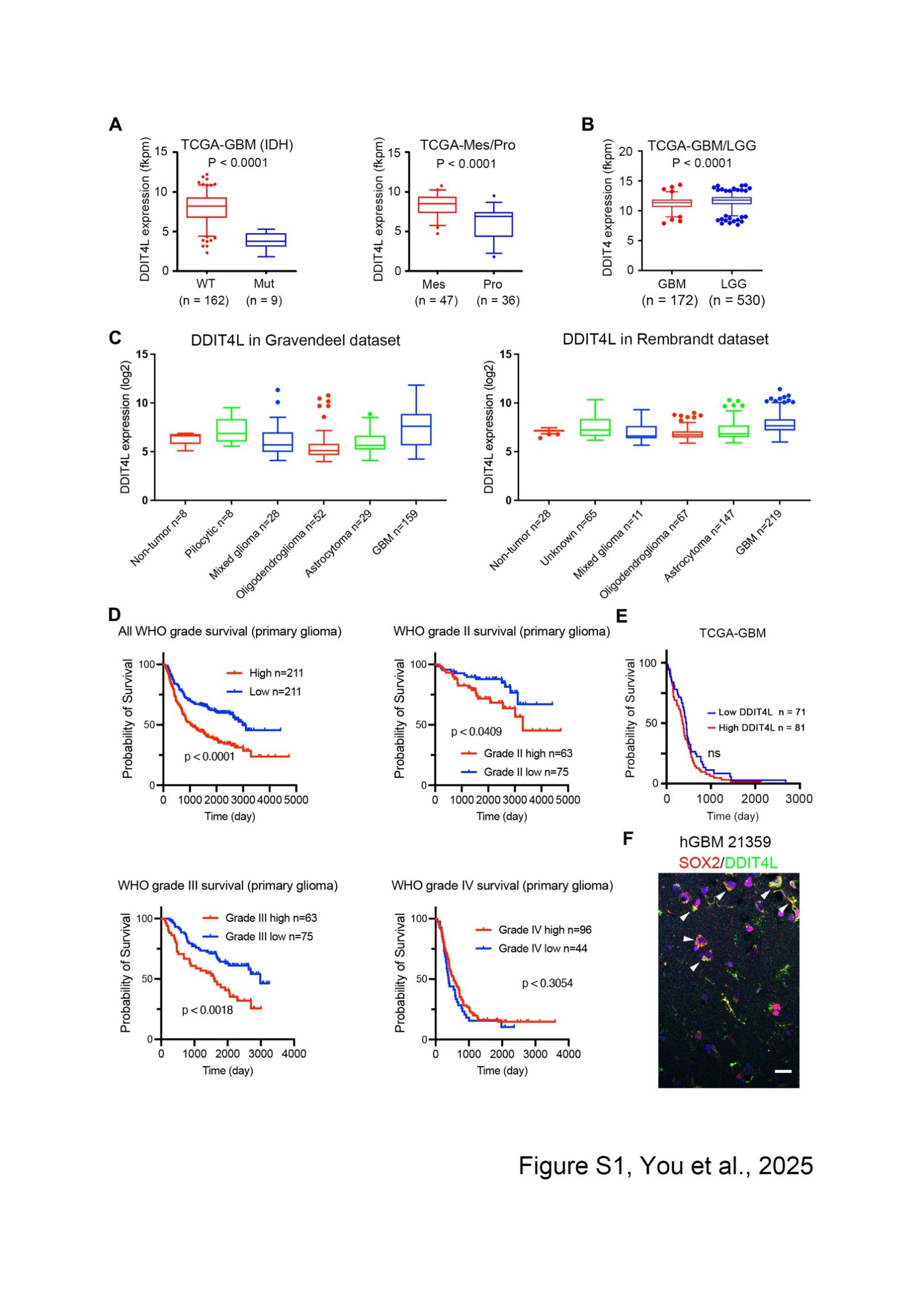

**Figure S1**. **DDIT4 and DDIT4L expression in GBM and LGG or in different types of glioma tumor tissues, Kaplan-Meier survival analysis in diffenrent WHO grades of glioma by the DDIT4L expression levels**

A. Boxplot of the DDIT4L expression in IDH wild-type & mutation GBM, and mesenchymal & proneural type of GBM from the TCGA database.
B. Boxplot of the *DDIT4* expression in GBM & LGG from the TCGA database. Significance testing was done with Mann-Whitney *U* test.
C. DDIT4L expression level in Gravendeel dataset and Rembrandt dataset were obtained from the online tool GlioVis (Visualization Tools for Glioma Datasets).
D. The CGGA database demonstrated the distinct survival time of glioma patients with different grades. Kaplan-Meier survival analysis of patients with different WHO grades of glioma stratified by the DDIT4L expression. Median DDIT4L expression was used for the stratification into DDIT4L-high and DDIT4L-low tumors.
E. The TCGA database demonstrated the similar survival time of GBM patients with different expression level of DDIT4L. Kaplan-Meier survival analysis of patients with GBM stratified by the DDIT4L expression. Median DDIT4L expression was used for the stratification into DDIT4L-high and DDIT4L-low tumors.
F. Double-immunostaining of DDIT4L and SOX2 in GBM showed DDIT4L co-localized with SOX2. Scale bar = 50 μm.

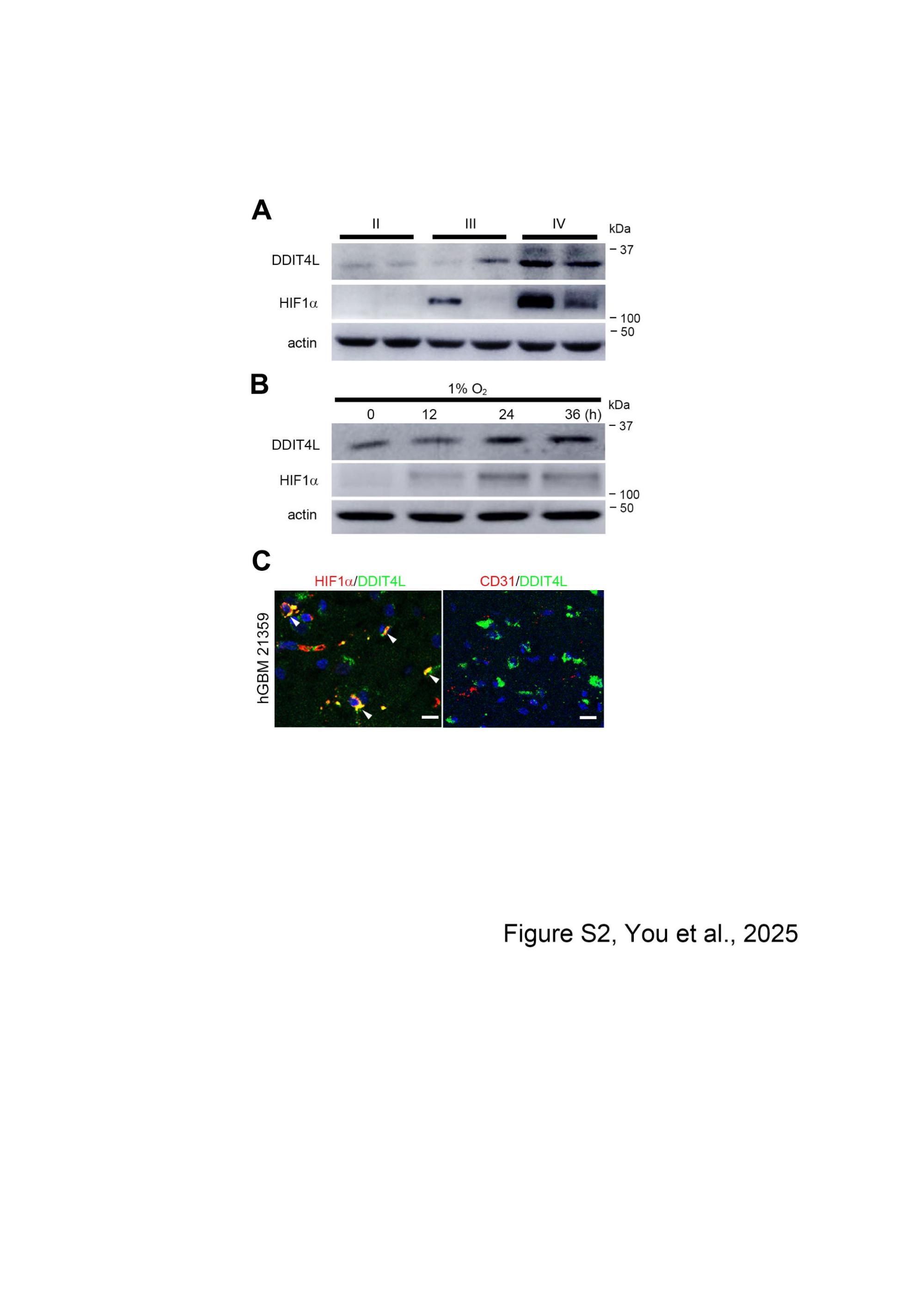

**Figure S2. Hypoxia up-regulates DDIT4L expression, DDIT4L Localizes in hypoxia-sensitive GBM cells**

**A, B**. DDIT4L and HIF1α expression increased with the aggression of glioma from grade II to grade IV (**A**), and increased in GBM cells exposure of hypoxia (1% O_2_) (**B**).

**C.** Double-immunostaining of DDIT4L and HIF1α, or DDIT4L and CD31 in GBM. Scale bar = 50 μm.

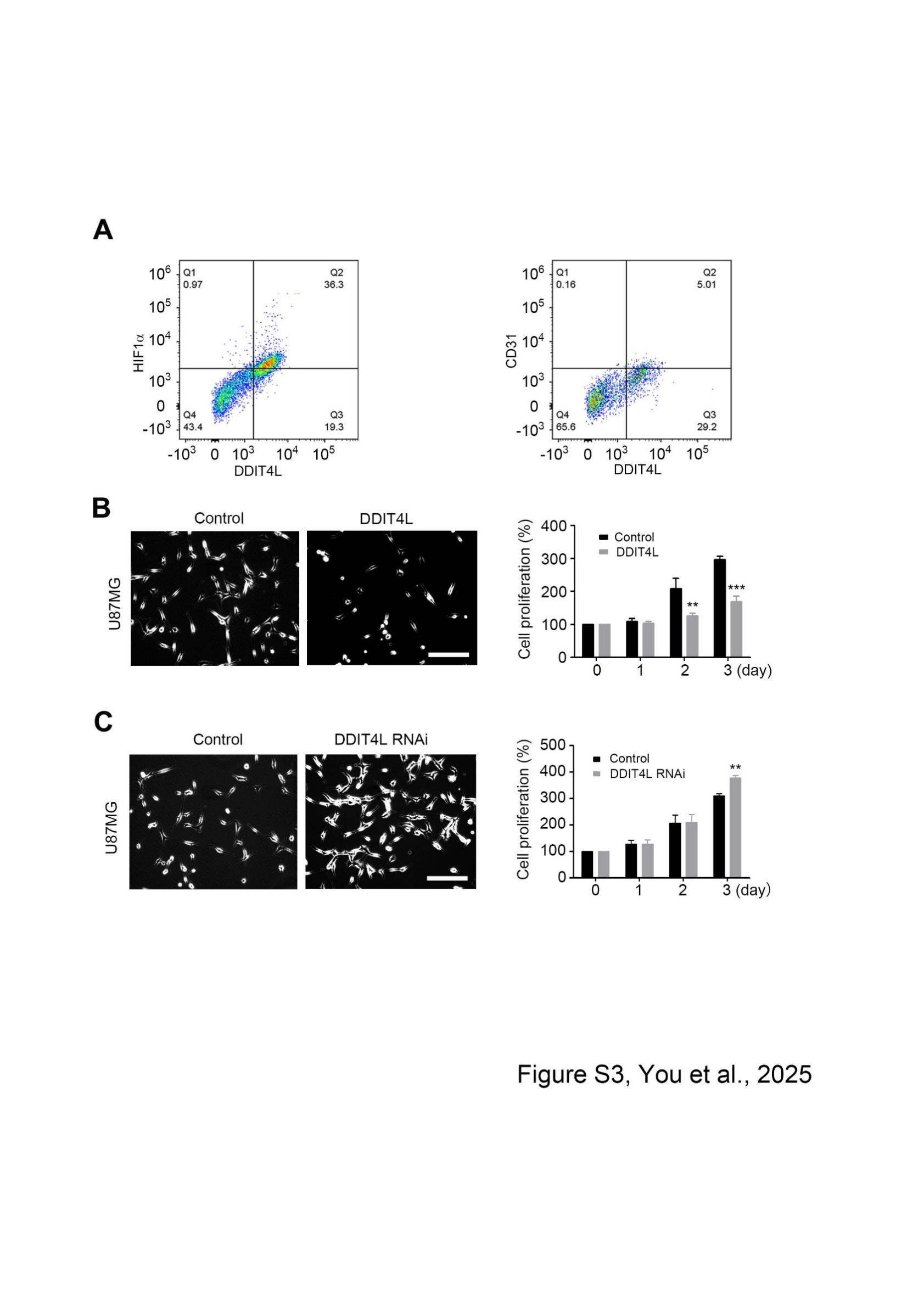

**Figure S3. DDIT4L suppresses GBM cells proliferation**

A. DDIT4L partially co-expressed with HIF1α, not with CD31 by the flow cytometry.
B. Transfected DDIT4L plasmid apparently inhibits U87MG cell proliferation 3 days after transfection.
C. Knockdown of DDIT4L by RNAi significantly enhances U87MG cell proliferation 3 days after transfection.

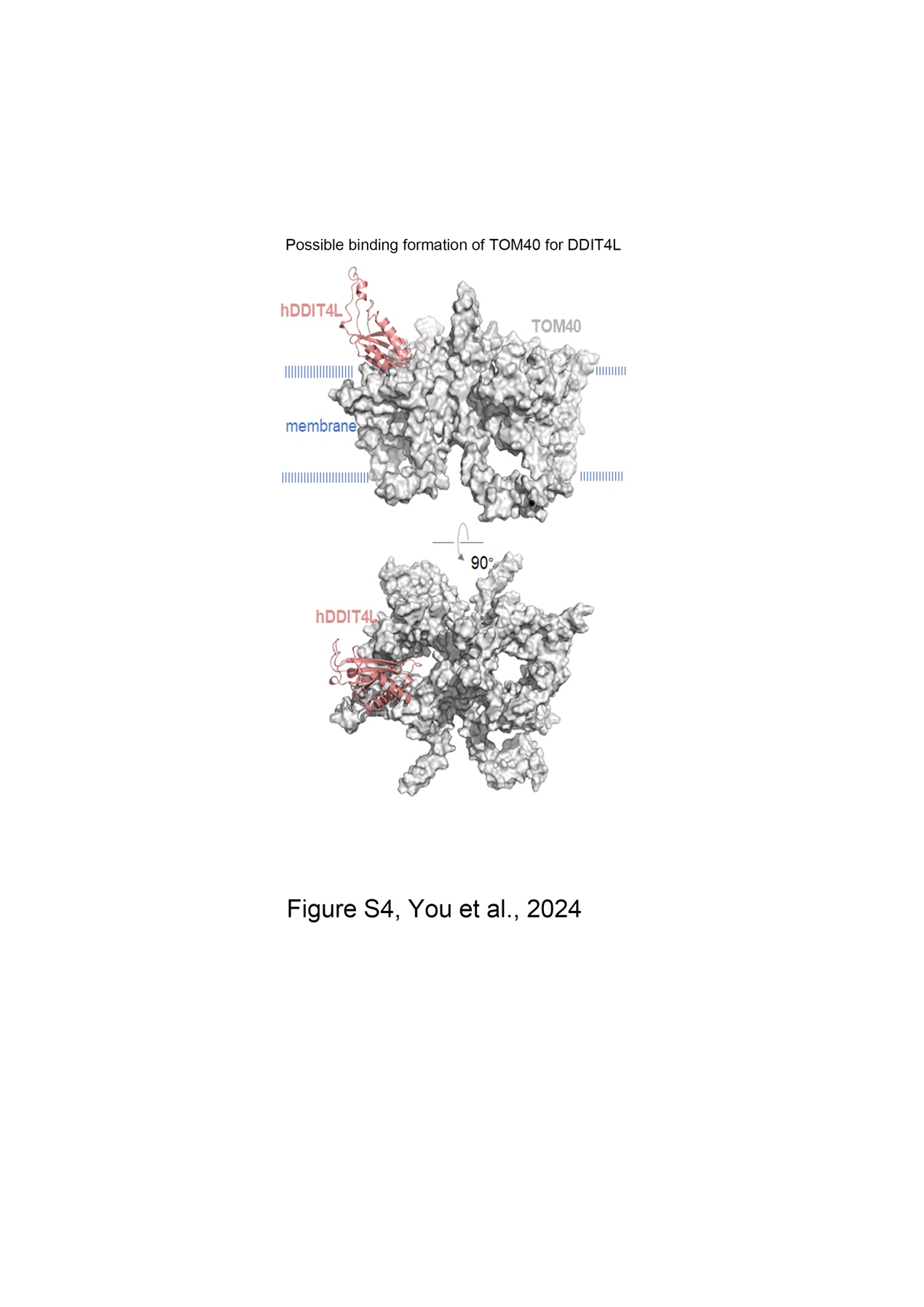

**Figure S4. Possible motifs implicated in interactions between DDIT4L and TOM40**

The selected DDIT4L (pink cartoon)/TOM40 (gray surface) model was derived from a cluster analysis of 2000 possible binding models based on ZDOCK scores and optimized by Zrank evaluation. The S96-C106P loop of DDIT4L, but not V125-P132 and K144-R106, might interact directly with TOM40.

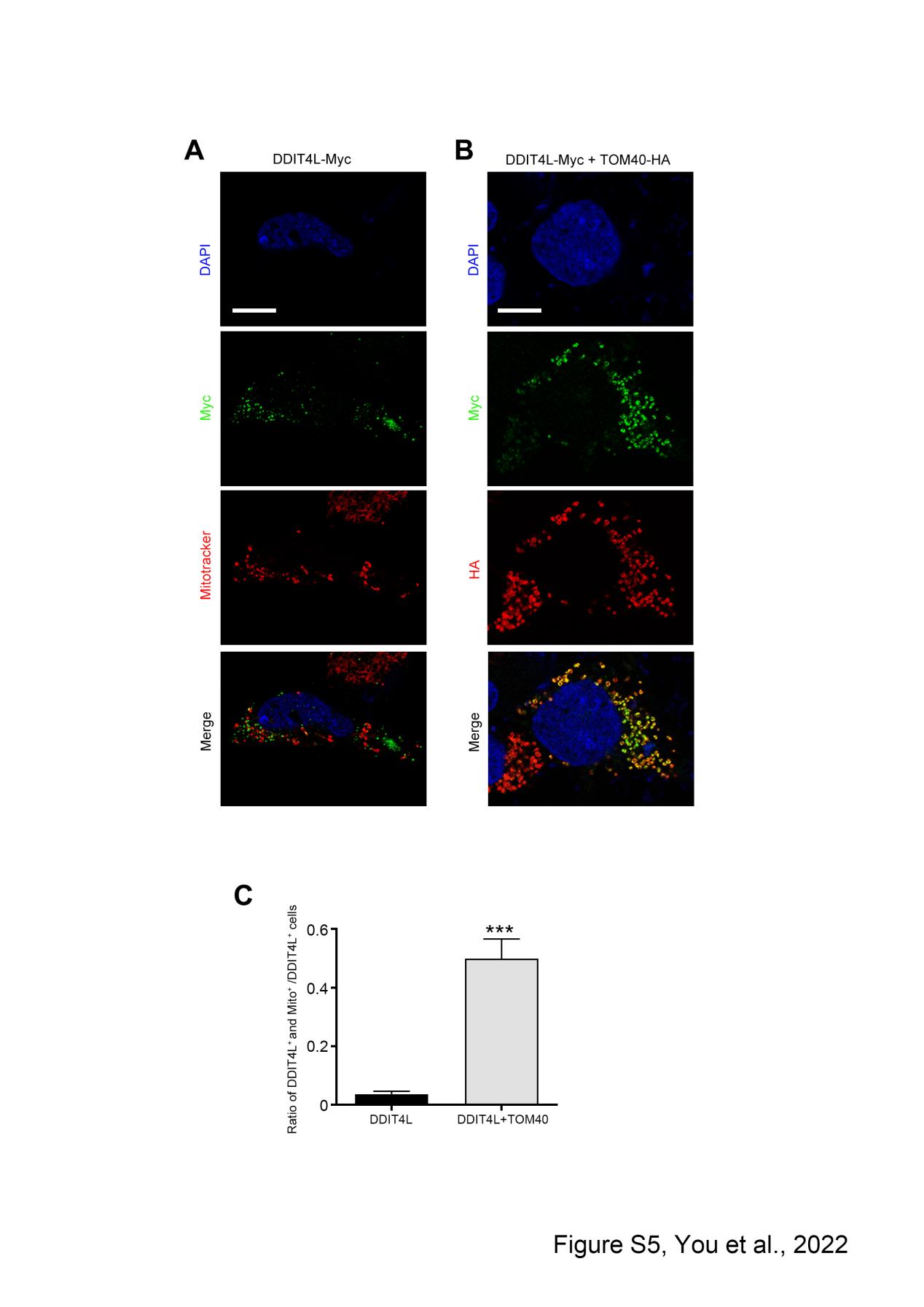

**Figure S5. Subcellular distribution of DDIT4L in HEK293 cells**

A. DDIT4L did not co-localize with mitochondria marked by mitochondria tracker (Mitotracker) after transfection of DDIT4L. Scale bar = 10 μm.
B. DDIT4L co-localized with mitochondria marked by TOM40 after co-transfection of DDIT4L and TOM40. Scale bar = 10 μm.
C. Quantification shows that the ratio of DDIT4L positive and Mitotracker positive cells/DDIT4L positive cells.

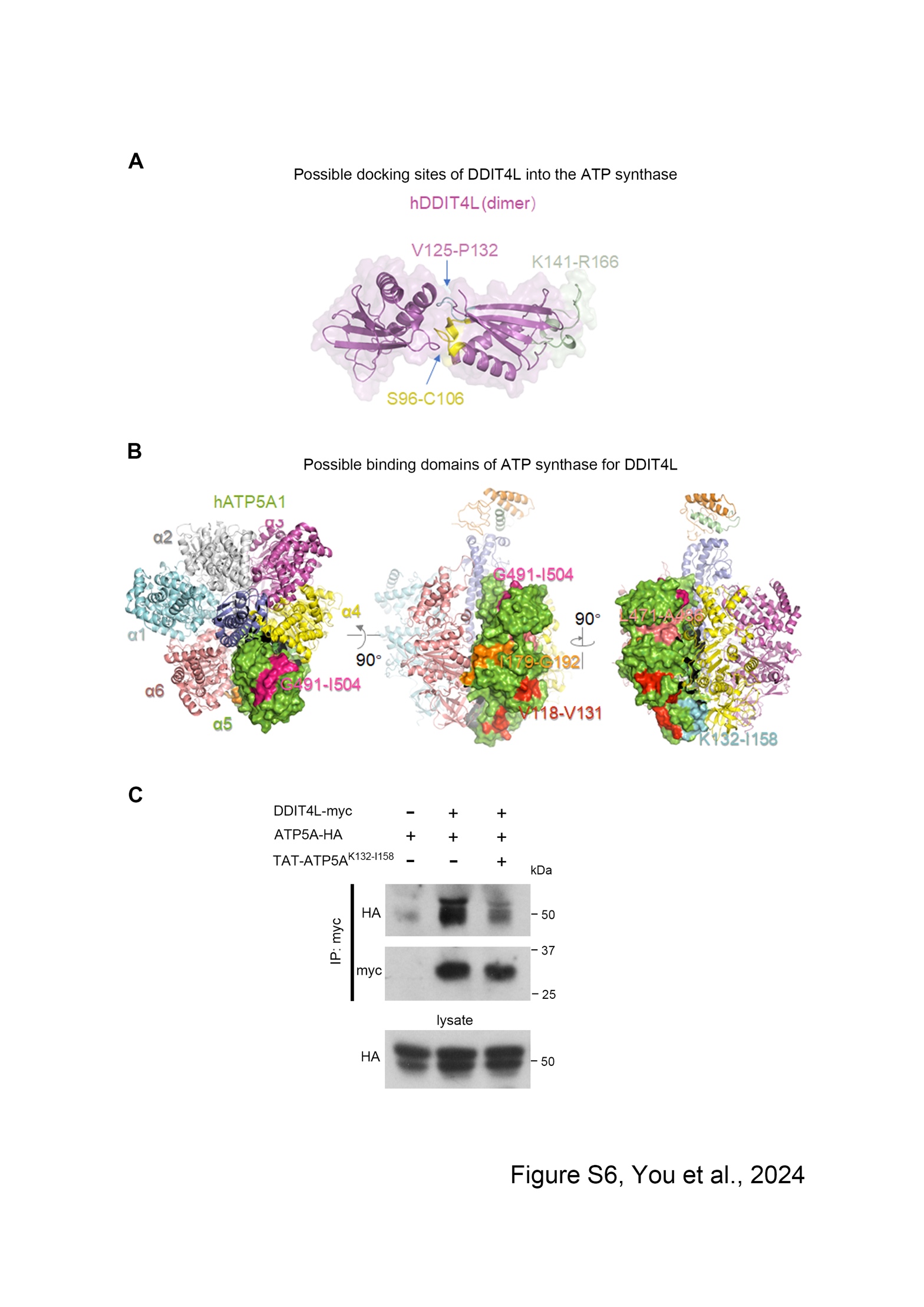

**Figure S6. Possible motifs implicated in interactions between DDIT4L and ATPase**

A. Possible docking site of DDIT4L in the ATPase using a combination of homology modeling and *in-silicon* docking. In this model, we screened out S96-C106, V125-P132 and K141-R166 motifs in DDIT4L as the possible binding domains for ATP5A.
B. Five motifs (V118-V131, K132-I158, I179-G192, L471-A486, G491-I504) of ATP5A had been predicted as the binding domains for DDIT4L using a combination of homology modeling and *in-silicon* docking.
C. The TAT-ATP5A^K132-I158^ peptide apparently reduced the interaction of DDIT4L and ATP5A in the transfected HEK293T cells (n = 3).

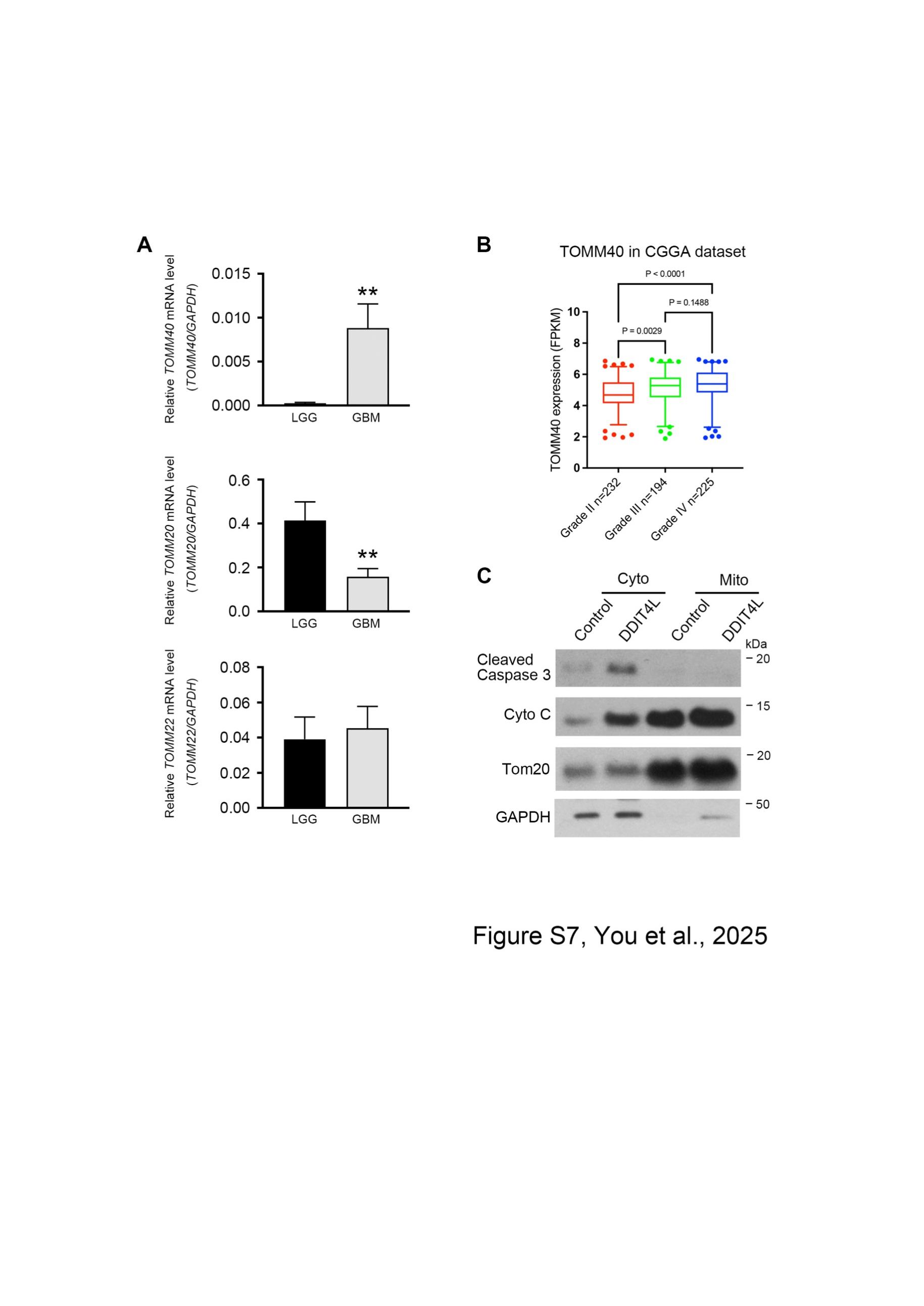

**Figure S7. TOM complex expression in GBM and DDIT4L** **inducing cell apoptosis in U87MG cells.**

A. The level of *TOMM40* mRNA was increased significantly, while level of *TOMM22* mRNA had no change and TOMM20 mRNA was decreased in GBM tissues.
B. The CGGA dataset demonstrated that TOMM40 expression level was increased in WHO grade III and IV when compared to that in Grade II gliomas.
C. Isolated mitochondria from cytoplasm in U87MG cells, the cytochrome c and cleaved caspase 3 significantly increase in cytoplasm after transfected DDIT4L (n = 3).

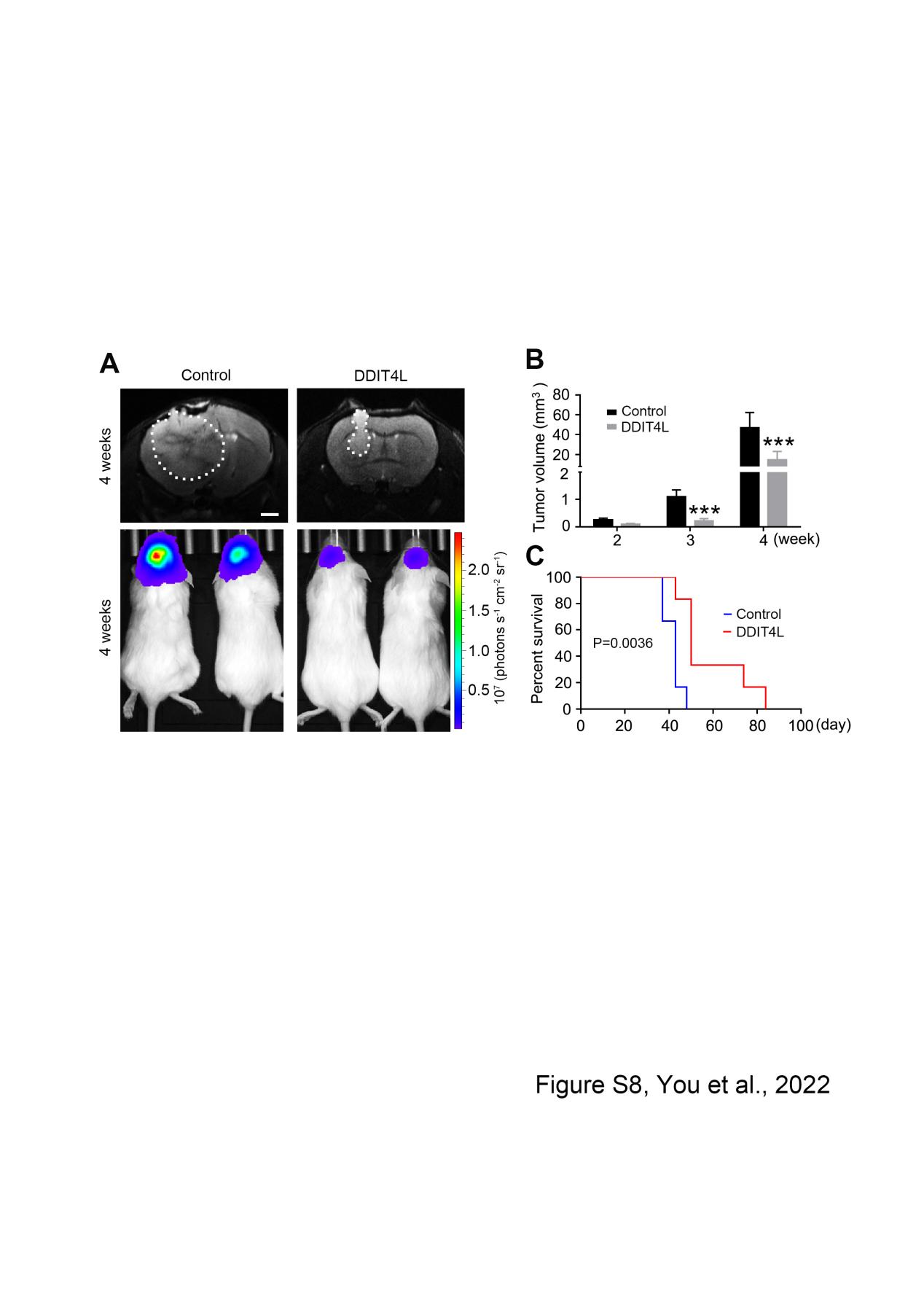

**Figure S8. DDIT4L suppresses tumor in the U87MG-derived model.**

A. MRI and bioluminescence imaging of GBM xenografts derived from U87MG cells expressing luciferase and with vector control or DDIT4L. Images from animals on day 28 after cells intracranial transplantation are shown. Scale bar = 1 mm.
B. Transfected DDIT4L apparently decreases tumor volume of GBM xenografts derived from U87MG cells (n = 6).
C. Kaplan-Meier survival analysis of mice with GBM xenografts after transfected DDIT4L.
